# Non-monotonic regulation of gene expression, neural progenitor fate and brain growth by the chromatin remodeller CHD8

**DOI:** 10.1101/469031

**Authors:** Shaun Hurley, Conor Mohan, Philipp Suetterlin, Jacob Ellegood, Fabrizio Rudari, Jason P. Lerch, Cathy Fernandes, M. Albert Basson

**Affiliations:** Centre for Craniofacial Regenerative Biology, King’s College London, London, United Kingdom; Department of Medical Biophysics, University of Toronto, Mouse Imaging Centre, Hospital for Sick Children, Toronto, Ontario, Canada; MRC Social, Genetic & Developmental Psychiatry Centre, Institute of Psychiatry, Psychology & Neuroscience, King’s College London, London, United Kingdom; MRC Centre for Neurodevelopmental Disorders, King’s College London, London, United Kingdom

**Author notes:** Corresponding author, (MAB). These authors contributed equally to this work (Shaun Hurley, Conor Mohan, Philipp Suetterlin).

**Keywords:** CHD8, autism, cortex, hypomorph, conditional knockout, mouse, intermediate progenitor, p53, gene expression, proliferation

## Abstract

Heterozygous *CHD8* mutations are associated with autism and macrocephaly with high penetrance in the human population. The reported mutations may have loss-of-function (haploinsufficient), hypomorphic or dominant negative effects on protein function. To determine the effects of reducing CHD8 protein function below haploinsufficient levels on brain development, we established a *Chd8* allelic series in the mouse. *Chd8* heterozygous mice exhibited relatively subtle brain overgrowth and little gene expression changes in the embryonic neocortex. In comparison, mild *Chd8* hypomorphs displayed significant postnatal lethality, with surviving animals exhibiting more pronounced brain hyperplasia, and significantly altered expression of over 2000 genes. Autism-associated genes were downregulated and neural progenitor proliferation genes upregulated. Severe *Chd8* hypomorphs displayed even greater transcriptional dysregulation, affecting genes and pathways that largely overlapped with those dysregulated in the mild hypomorphs. By contrast, homozygous, conditional deletion of *Chd8* in early neuronal progenitors resulted in the induction of p53 target genes, cell cycle exit, apoptosis and pronounced brain hypoplasia. Intriguingly, increased progenitor proliferation in hypomorphs was primarily restricted to TBR2+ intermediate progenitors, suggesting critical roles for CHD8 in regulating the expansion of this population. Given the importance of these progenitors in human cortical growth, this observation suggests that human brain development might be more sensitive to CHD8 deficiency than the mouse. We conclude that brain development is acutely sensitive to CHD8 dosage and that the varying sensitivities of different progenitor populations and cellular processes to CHD8 dosage can result in non-linear effects on gene transcription and brain growth.

## Introduction

Developments in exome sequencing have recently facilitated the discovery of de novo likely gene disrupting mutations in *CHD8* (Chromodomain helicase DNA-binding protein 8) in individuals with autism spectrum disorder (ASD) (Iossifov et al., 2014; Neale et al., 2012; O’Roak et al., 2012a; O’Roak et al., 2012b; Talkowski et al., 2012). Mutations in *CHD8* remain one of the highest confidence ASD risk factors identified to date and individuals with *CHD8* mutations exhibit autism (96%), macrocephaly (64%), craniofacial abnormalities (78%) and anxiety (27%) (Bernier et al., 2014; Stessman et al., 2017). *CHD8* encodes a member of the ATP-dependent CHD chromatin remodelling family (Thompson et al., 2008).

CHD8 was initially identified as a direct repressor of β-catenin and p53 target genes (Nishiyama et al., 2004; Nishiyama et al., 2009; Nishiyama et al., 2012; Sakamoto et al., 2000). Early embryonic lethality of homozygous *Chd8* deletion in the mouse is associated with p53-mediated apoptosis, consistent with its role as a transcriptional repressor of p53 target genes (Nishiyama et al., 2004). CHD8 appears to function as a positive regulator of other ASD-associated genes in human neural progenitor cells (Cotney et al., 2015; Sugathan et al., 2014), implicating reduced expression of these genes in neural progenitors as a potential mechanism underlying neurodevelopmental phenotypes. Evidence for mild brain overgrowth, reminiscent of the macrocephaly observed in patients with *CHD8* mutations, has been reported recently in several different *Chd8* heterozygous mouse models (Gompers et al., 2017; Katayama et al., 2016; Platt et al., 2017; Suetterlin et al., 2018).

To explore the transcriptional dysregulation that may underlie abnormal brain development in heterozygous mice, gene expression has been investigated at different stages of brain development. These studies have revealed subtle gene expression changes in *Chd8*^+/-^ mice during embryonic development (Gompers et al., 2017; Suetterlin et al., 2018). By contrast, in vitro studies on neural progenitor cells have identified more substantial transcriptional dysregulation arising from CHD8 knock-down. Sugathan et al. observed 1756 differentially expressed genes (DEGs) upon *CHD8* knock-down in human iPSC-derived neural progenitor cells (Sugathan et al., 2014). Both this and another study have demonstrated that CHD8 is typically recruited to promoters enriched for transcriptionally-permissive chromatin marks, suggesting a role for CHD8 in transcriptional activation (Cotney et al., 2015; Sugathan et al., 2014).

The striking phenotypes in human individuals with *CHD8* mutations and pronounced gene expression changes in neural progenitor cell lines, contrast with the mild brain and embryonic transcriptional abnormalities observed so far in *Chd8* heterozygous mice. The only study so far to report convincing ASD-like behavioural phenotypes associated with *Chd8* deficiency, involved *Chd8* knock-down to ~20% of wildtype protein levels by in utero electroporation of upper layer neuronal progenitors (Durak et al., 2016). Together, these findings suggest that *CHD8* haploinsufficiency has more pronounced, or unique effects on human brain development, or that some human mutations reduce CHD8 function by more than 50%.

To examine the possibility that certain phenotypes may only appear at sub-heterozygous *Chd8* levels in the mouse, we created an allelic series of *Chd8*-deficient mice to reduce CHD8 protein step-wise to approximately 25% (mild hypomorph), 12% (severe hypomorph) and 0% (conditional knockout) of wildtype levels. Mild hypomorphs exhibited more pronounced brain overgrowth than heterozygous mice. We identified a distinct role for CHD8 in regulating the proliferation of TBR2+ intermediate progenitors. As this cell type is responsible for increased human brain expansion during evolution, this finding suggests that CHD8 may have more pronounced, or even unique effects on human brain development. Gene expression analyses of hypomorphic mice identified many important neurodevelopmental, autism-associated and cell cycle genes that showed a non-graded threshold transcriptional response as CHD8 levels fell below those in heterozygous mice. Analysis of conditional knockout mice revealed an additional threshold transcriptional response of p53-reguated genes, and non-monotonic effects on cell cycle genes, leading to pronounced cortical hypoplasia, in contrast to the cortical hyperplasia in heterozygous and mild hypomorphic mice. These observations suggest that mutations and other factors that reduce CHD8 function below 50% may have disproportionally large, and unexpected effects on gene expression and brain development.

## Results

### *Chd8* hypomorphs exhibit reduced survival and exacerbated brain overgrowth

We generated a hypomorphic mouse *Chd8* allele (*Chd8^neo^*, Fig. 1A) by inserting a neo cassette between exons 3 and 4 to reduce gene expression through splicing and termination of transcripts (Meyers et al., 1998). Aberrant splicing of *Chd8* transcripts into the neo cassette was observed (Fig. 1A). *Chd8^neo/neo^* and *Chd8*^*neo/*-^ embryos showed 85% and 88% reductions in *Chd8* transcripts, respectively, compared to the 63% decrease in *Chd8*^+/-^ embryos (Fig. 1B). Full-length CHD8 protein levels were reduced by 49% in *Chd8*^+/-^, 73% in *Chd8^neo/neo^*, and 87% in *Chd8*^*neo/*-^ neocortices, with no evidence for remaining truncated CHD8 protein products (Fig. 1C, Suppl. Fig. 1).

**Fig 1.**
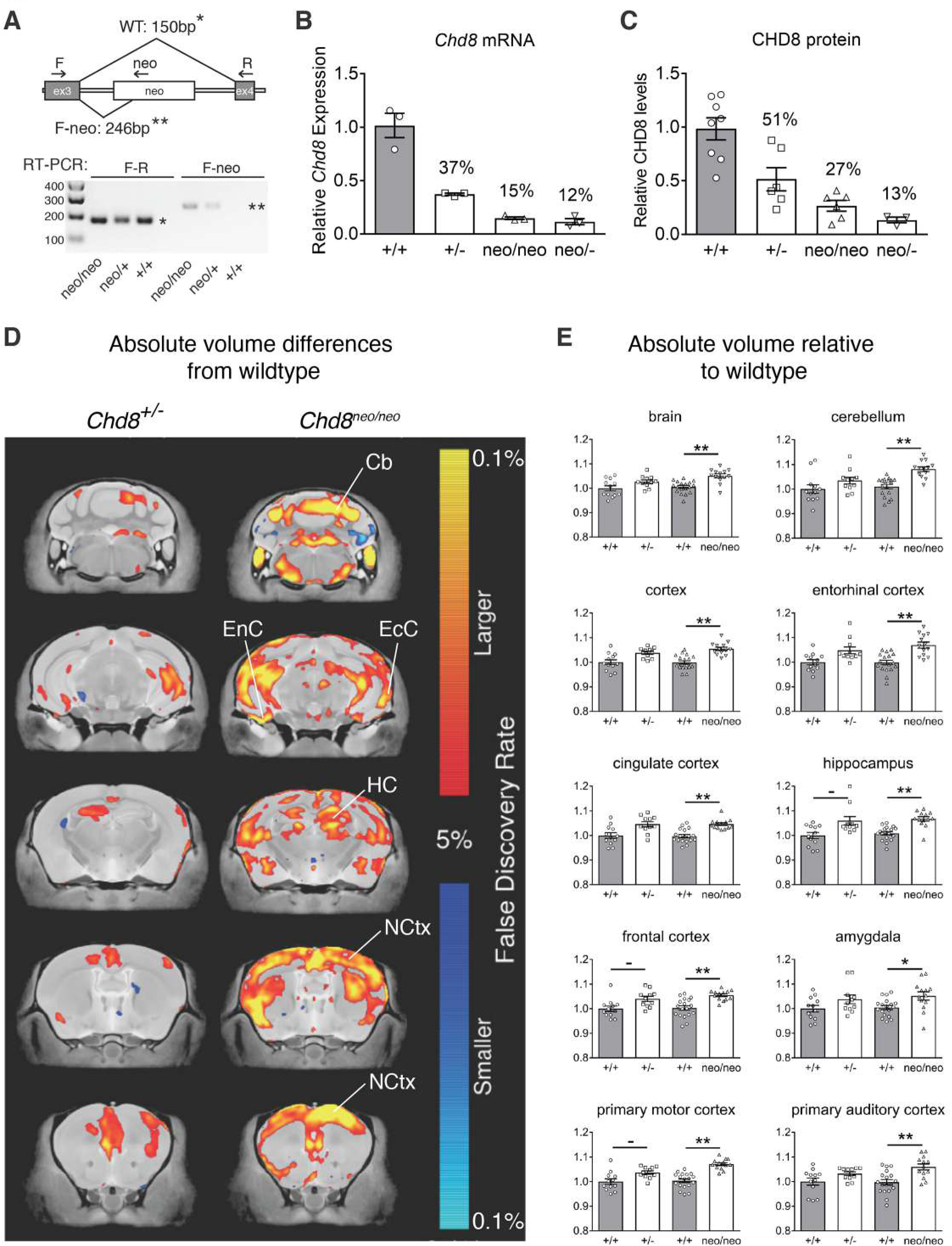
Volumetric brain changes in *Chd8*-deficient mice. A. Diagrammatic representation of the *Chd8* allele containing the neo cassette between exons 3 and 4. Exon 3 splicing to exon 4 yields a 150bp product (*) by RT-PCR using primers F andR. Aberrant splicing from exon 3 in to the neo cassette yields a 246bp product (**) with primers F and neo.
B. Quantitative RT-PCR of *Chd8* transcripts in E9.5-E10.5 neocortices of indicated genotypes.
C. Western blotting of CHD8 protein in E12.5 neocortices of indicated genotypes.
D. High-resolution 7T structural MRI coronal images of *Chd8*^+/-^ and *Chd8^neo/neo^* brains, from posterior (top) to anterior (bottom) are shown. Absolute volumetric differences, relative to wildtype controls are coloured according to the scale on the right. Some regions with enlarged volumes are labeled as follows: NCtx-neocortex, EcC – ectorhinal cortex, EnC – entorhinal cortex, HC – hippocampus, Cb – cerebellum.
E. Absolute volumes relative to wildtypes are plotted for whole brain, neocortex and several other brain regions for the different genotypes as indicated. ^-^FDR<0.15, *FDR<0.05, **FDR<0.01. See also Supplementary Table 1.

The additional reduction in CHD8 levels in *Chd8^neo/neo^* mice from heterozygotes led to a significant reduction in postnatal survival (Table 1). As CHD8 is expressed in multiple tissues during development (Kasah et al.), postnatal lethality is likely as a result of congenital defects in essential organs.

**Table 1:**
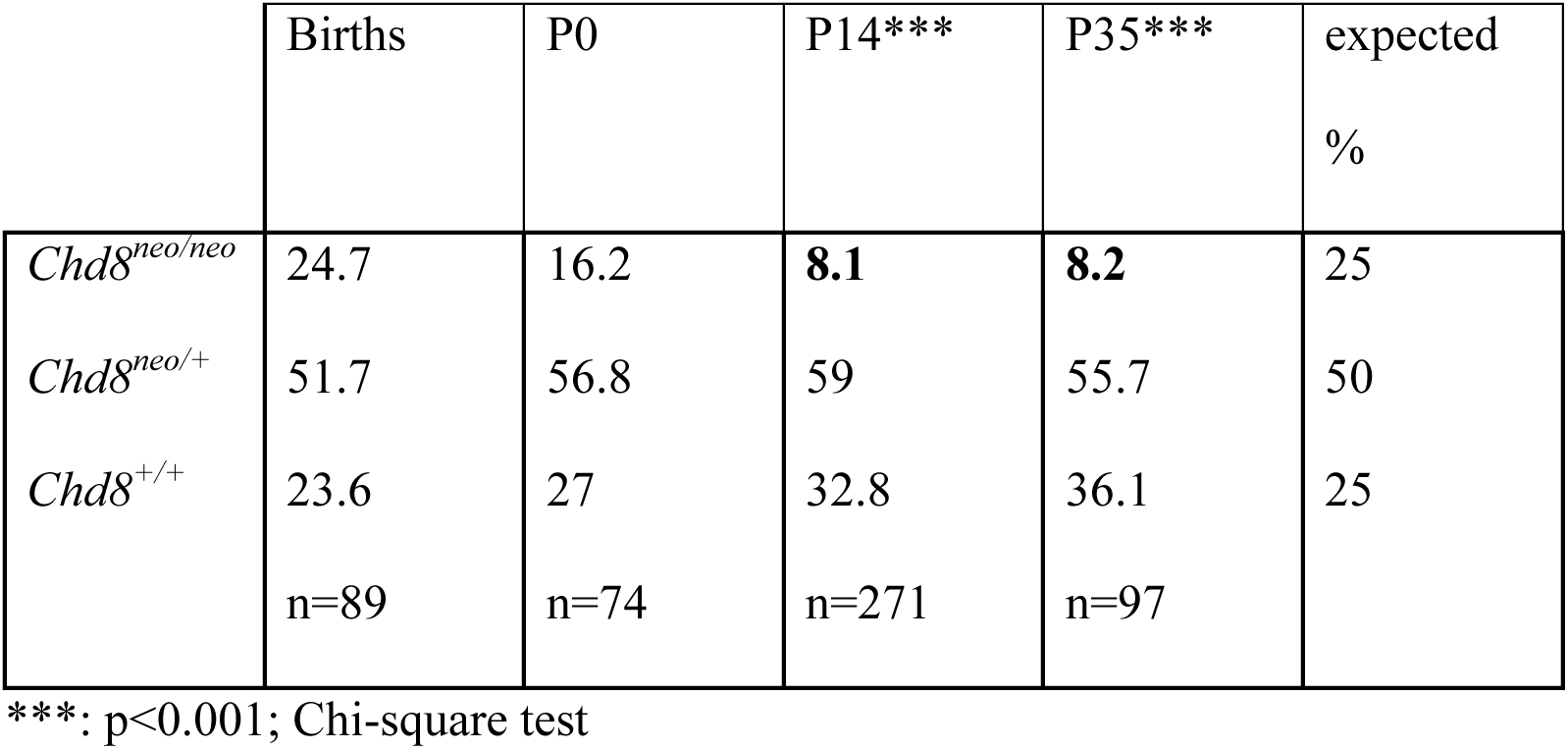
Reduced postnatal survival of *Chd8^neo/neo^* pups.

High resolution structural MRI revealed volumetric increases in a number of brain regions in *Chd8^neo/neo^* mice compared to wildtype littermates (Fig. 1D). This phenotype was more pronounced compared to *Chd8*^+/-^ mice (Fig. 1D), with total brain volume increased by 4.5% in *Chd8^neo/neo^* mice, compared to the 2.7% increase in *Chd8*^+/-^ mice (Fig. 1E). Several regions that showed evidence of overgrowth in *Chd8*^+/-^ mice (Fig. 1E, Table 2) demonstrated robust increases in volume in *Chd8^neo/neo^* mice, including the frontal, cingulate and entorhinal cortices and the hippocampus (Fig. 1E, Table 2, Suppl. Table 1).

**Table 2.** Brain volume differences relative to control wildtype littermates.

| Brain area | *Chd8* genotype |  |
| --- | --- | --- |
|  | +/- | neo/neo |
| brain volume | 2.73% | 4.53%** |
| cortex | 3.82% - | 5.53%** |
| cerebellum | 3.41% | 6.99%** |
| hippocampus | 5.97% - | 5.84%** |
| primary motor cortex | 3.68% - | 6.68%** |
| primary somatosensory cortex | 3.7% - | 5.62%** |
| primary auditory cortex | 3.2% | 6.23%** |
| primary visual cortex | 3.39% | 7.15%** |
| frontal cortex | 3.99% - | 5.21%** |
| frontal association cortex | 4.81% - | 6.07%** |
| entorhinal cortex | 4.83% - | 7.05%** |
| cingulate cortex | 4.57% - | 4.84%** |
| amygdala | 3.82% | 4.64%* |

-FDR<0.15; \* FDR<0.05; \*\*FDR<0.01

Mice were born at expected Mendelian frequencies from *Chd8*^*neo/*+^ intercrosses. Percentage survival is shown for the different genotypes at P0, P14 and P35. Note the significantly reduced observed percentage of homozygous *Chd8^neo/neo^* mutants at P14 and P35 (***p<0.001, Chi-square test) in two independent groups of mice.

### Stepwise reductions in CHD8 levels result in progressively more pronounced gene expression changes

To determine the impact of sub-heterozygous *Chd8* levels on gene expression in the E12.5 neocortex, RNA-sequencing (RNA-seq) was performed. Data from heterozygous (*Chd8*^+/-^), mild (*Chd8^neo/neo^*) and severe (*Chd8*^*neo/*-^) hypomorphs, together with their respective wildtype littermate controls were included for analysis.

This analysis identified 14 differentially expressed genes (DEGs, excluding *Chd8*, FDR <0.05) in *Chd8*^+/-^ (Fig. 2A, Suppl. Table 2) and 2209 DEGs in *Chd8^neo/neo^* neocortices (Fig. 2B, Suppl. Table 2), indicating that many CHD8-regulated genes only show significant transcriptional effects when CHD8 levels fall below 50%. In *Chd8*^neo/-^ embryos, 2592 DEGs (FDR <0.05) were identified (Fig. 2C, Suppl. Table 2). The visualization of differential gene expression in a heat map demonstrated the marked transcriptomic differences between heterozygotes and mild hypomorphs (Fig. 2D). DEGs could be divided into four groups based on their responses to reduced CHD8 levels: 1) genes that show a linear response to *Chd8* downregulation (e.g. *Tet1* and *Zcwpw1*, Fig. 2E), 2) genes that are not significantly different in *Chd8*^+/-^ embryos but sharply up- or downregulated in *Chd8^neo/neo^* embryos (e.g. *Nlgn3* and *Slc1a5*, Fig. 2E), 3) genes that are only significantly dysregulated in *Chd8*^*neo/*-^ embryos (e.g. *Gpat2*), and 4) genes that exhibited non-linear responses (e.g. *Slc9b2*, Fig. 2E). The majority of DEGs (>99%) fell within group 2, in agreement with the finding that over 2000 genes showed a striking threshold response as CHD8 protein is reduced by 24% from heterozygotes to mild hypomorphs.

**Fig 2.**
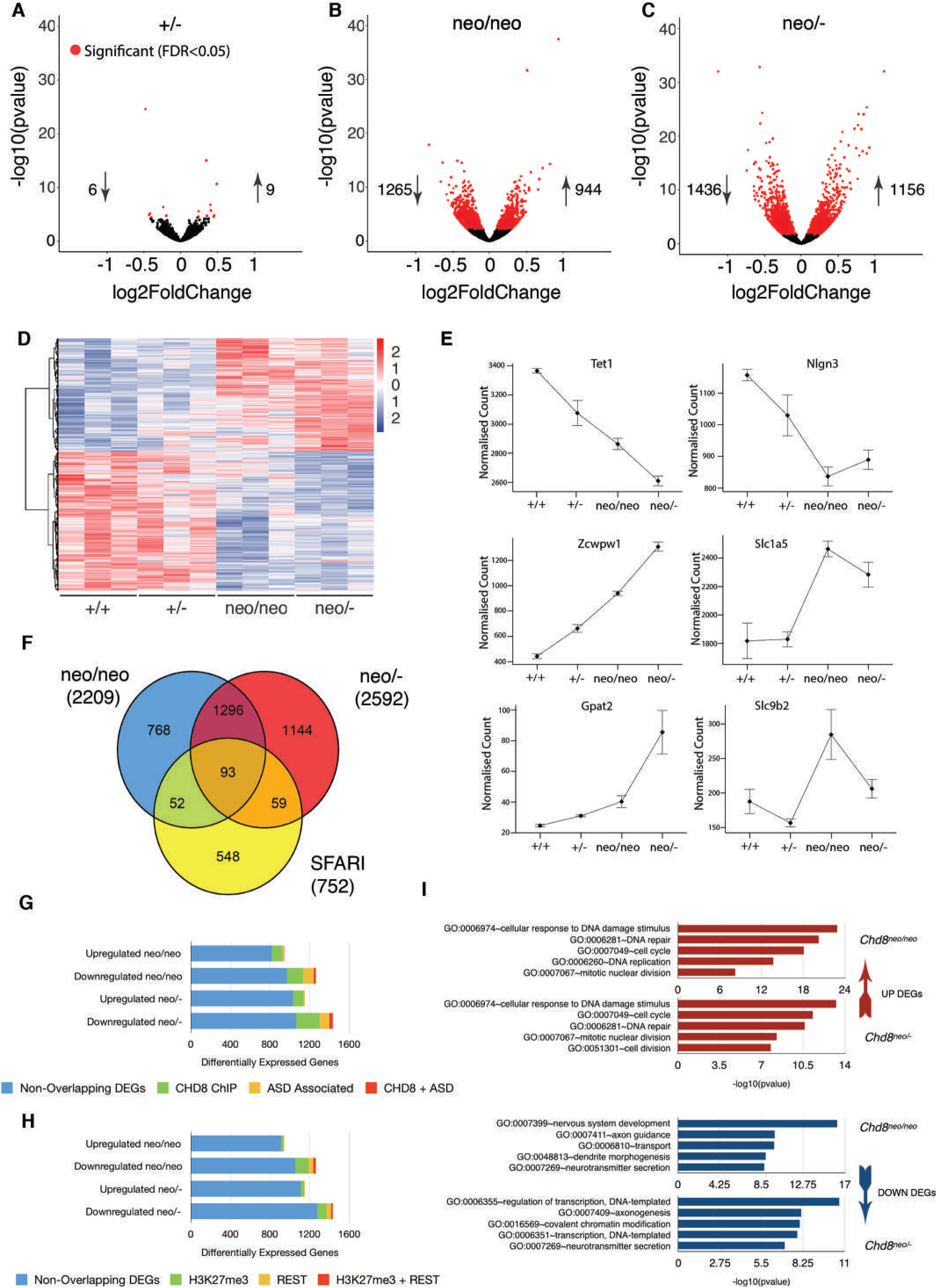
Gene expression changes in *Chd8*-deficient neocortices. A. Volcano plot indicating all DEGs detected by RNA-seq in E12.5 *Chd8*^+/-^ embryonic cortices. Each point represents an individual gene, and all heterozygote DEGs (FDR <0.05) are highlighted in red.
B. Volcano plot showing all DEGs in E12.5 neo/neo cortex. All differentially expressed genes (FDR <0.05) are highlighted in red.
C. Volcano plot of all DEGs in E12.5 neo/- cortex. All differentially expressed genes (FDR <0.05) are highlighted in red.
D. Heatmap of genes differentially expressed in neo/neo and neo/- embryos, indicating transformed relative expression levels in +/+, +/-, neo/neo and neo/- embryos.
E. Mean normalised count of aligned RNA-seq reads for a selection of genes that were differentially expressed in the mild and severe hypomorphs.
F. Venn diagram showing extent of overlap between neo/neo and neo/- DEGs and ASD associated genes obtained from the SFARI Human Gene database.
G. Breakdown of neo/neo and neo/- DEGs that are ASD associated, DEGs with known CHD8 binding sites, and DEGs that are both ASD associated and have known CHD8 binding sites.
H. Breakdown of neo/neo and neo/- DEGs that are decorated with H3K27me3 in neural progenitors, DEGs with known REST binding sites, and DEGs that have been shown to have both H3K27me3 in neural progenitors and REST binding sites.
I. Gene Ontology (GO) analysis of up and down-regulated neo/neo and neo/- DEGs under the “Biological Processes” category. The five most significant hits are shown for each set. See also Supplementary Table 2.

Comparing DEGs in *Chd8^neo/neo^* and *Chd8*^*neo/*-^ samples identified 1389 genes common to both datasets (Fig. 2F), all of which were changed in the same direction. ASD-associated genes were highly enriched in the DEGs from both *Chd8^neo/neo^* (145 genes, p=1.32 × 10-9, OR = 1.83, Fisher’s exact test for count data, Fig. 2F, Suppl. Table 2) and *Chd8*^*neo/*-^ embryos (152 genes, p= 5.532 x10-7, OR = 1.62, Fisher’s exact test for count data, Suppl. Table 2). Nearly half (46%) of these ASD-associated genes were common to *Chd8^neo/neo^* and *Chd8*^*neo/*-^ mice (Fig. 2F). The majority (89% and 88%, respectively) of ASD-associated DEGs were downregulated (Fig.3G, Suppl. Table 2), supporting the idea of CHD8 as an important positive regulator of neurodevelopmental genes (Cotney et al., 2015; Sugathan et al., 2014).

**Fig 3.**
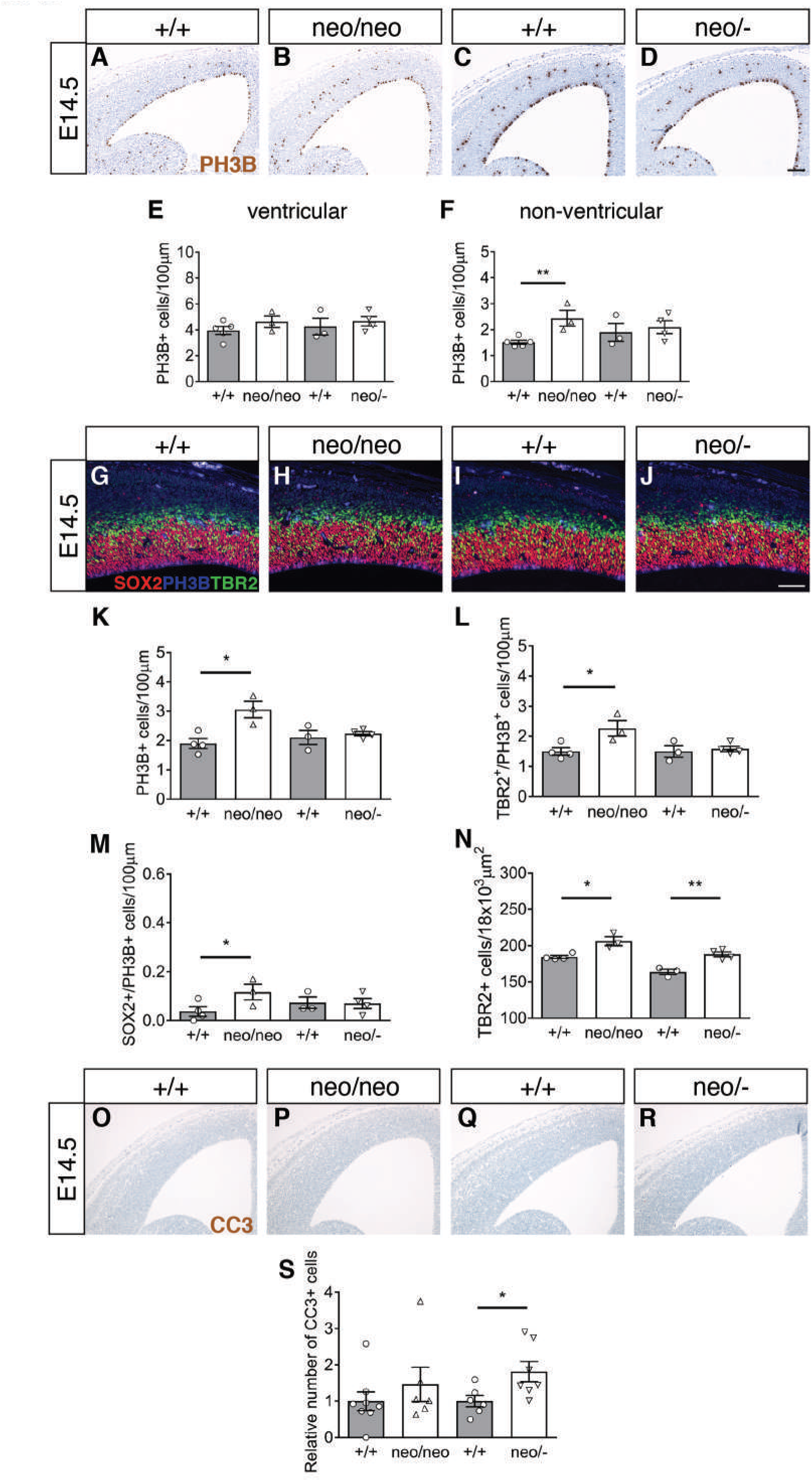
Increased proliferation of basal neural progenitors in *Chd8^neo/neo^* embryos. A,B,C,D) Immunohistochemistry to detect PH3B+ nuclei (brown) in coronal sections through the telencephalon of E14.5 embryos. Scale bar = 100μm. E,F) Quantification of PH3B+ cells per 100μm of neocortex in either ventricular (E) or non-ventricular (F) areas in E14.5 embryos (+/+, n=5; neo/neo, n=3; +/+, n=3; neo/-, n= 4) Mean±SEM, **p<0.005, student’s t-test between *Chd8* mutant and respective wildtype littermates). G,H,I,J) Immunostaining for SOX2 (red), PH3B (blue) and TBR2 (green) in E14.5 embryo neocortex. Scale bar = 50μm. K,L,M) Quantification of non-ventricular PH3B+ (K), TBR2+/PH3B+ (L) and SOX2+/PH3B+ (M) cells per 100μm of neocortex in E14.5 embryos (+/+, n=4; neo/neo, n=3; +/+, n=3; neo/-, n= 4) Mean±SEM, *p<0.05, student’s t-test between *Chd8* mutant and respective wildtype littermates. N) Quantification of subventricular TBR2+/SOX2-cells per 1800μm^2^ in E14.5 embryos (+/+,n=4; neo/neo, n=3; +/+, n=3; neo/-, n= 4) Mean±SEM, *p<0.05, **p<0.01, student’s t-test between *Chd8* mutant and respective wildtype littermates). O,P,Q,R) Cleaved caspase 3 (CC3) immunostaining of dorsal neocortex of E14.5 embryos. Scale bar = 100μm. S) Quantification of CC3+ cells per 100μm of neocortex in E14.5 *Chd8* mutant embryos shown normalised to respective wildtype littermates (+/+, n=4; neo/neo, n=3; +/+, n=3; neo/-, n=3; Mean±SEM).

To identify genes possibly directly regulated by CHD8, ChIP-seq data from Cotney et al. was used to identify gene promoters with CHD8 recruitment in embryonic mouse brain, human brain and neural progenitors (Cotney et al., 2015). A relatively small fraction (14% *Chd8^neo/neo^* and 15% *Chd8*^*neo/*-^) of DEGs appear to be directly regulated by CHD8 (Fig. 2G, Suppl. Table 2), consistent with evidence that CHD8 regulates the expression of many other chromatin and epigenetic modifiers (Cotney et al., 2015; Sugathan et al., 2014). CHD8 was present at both up- and downregulated genes (Fig. 2G, Suppl. Table 2), suggesting that CHD8 can function as both an activator and repressor of genes during cortical development.

To identify potential transcriptional co-regulators and DNA-binding factors that may cooperate with CHD8 during embryonic cortical development, Gene Set Enrichment Analysis was performed using the “ENCODE and ChEA Consensus TFs from ChIP-X” database in Enrichr (Chen et al., 2013). This analysis revealed an over-representation of E2F (E2F4, E2F6 and E2F1) targets in the upregulated genes (Suppl. Fig. 2, Suppl. Table 2). E2Fs compose a family of transcription factors with important roles in DNA replication, cell cycle progression and proliferation. CHD8 has been previously shown to be involved in E2F-dependent transcriptional activation, and is necessary for recruitment of the "activator" E2F transcription factors E2F1 and E2F3 to G1/S transition promoters (Subtil-Rodriguez et al., 2014). Our findings suggest that CHD8 functions as a repressor of E2F-regulated genes in the developing cortex and implicate increased progenitor proliferation as a potential mechanism for the brain hyperplasia in these mice.

For downregulated genes, an over-representation of targets of REST (RE1-Silencing Transcription factor) and the Polycomb component Suz12 was seen (Suppl. Fig. 2, Suppl. Table 2). As Suz12 is essential for the activity and stability of the PRC2 complex, we asked if any of the DEGs are marked by the PRC2-repressive modification H3K27me3 in normal neural progenitor cells (Mohn et al., 2008). The majority of DEGs that were marked by H3K27me3 in neural progenitors were downregulated in hypomorphic mice (Fig. 2H), suggesting that CHD8 may indeed regulate their expression by antagonising Polycomb repression. REST is a master regulator of neurodevelopment. REST has been shown to directly interact with CHD8, and is abnormally activated in *Chd8* haploinsufficient mouse brain (Katayama et al., 2016). Overlapping DEGs with REST ChIP-seq data (Johnson et al., 2008), found REST target genes predominantly amongst downregulated genes (Fig. 2H), supporting the notion that aberrant REST activation in *Chd8*-deficient embryonic brain may contribute to gene repression. Furthermore, 40% of the downregulated REST target genes are also marked by H3K27me3 in neural progenitor cells, implying roles for both REST and Polycomb in the repression of these genes in *Chd8* hypomorphs (Fig. 2H).

Gene ontology analyses showed a significant enrichment of cell cycle, DNA replication and repair genes in the upregulated genes in hypomorphs (Fig. 2I, Suppl. Table 3). Neurodevelopmental gene categories were enriched in the doewnregulated gene sets (Fig. 2I, Suppl. Table 3).

### *Chd8* deficiency increases proliferation of cortical progenitors outside of the ventricular zone during mid-neurogenesis

To explore whether increased expression of cell cycle and DNA replication genes in hypomorphs may be associated with increased progenitor expansion during embryonic development, we immunolabelled progenitors in the G2/M phase of mitosis in coronal brain sections with an antibody against phosphohistone-3B (PH3B). No difference in the number of mitotic progenitors was observed in the E12.5 *Chd8^neo/neo^* neocortex compared to wildtype littermates (Suppl. Fig. 3A-C). To determine if differences may arise later, we performed the same experiment at E14.5 (Fig 3A-D). At this stage, the number of proliferating progenitors in the *Chd8^neo/neo^* neocortical ventricular zone showed a trend towards a small increase (Fig. 3E). Intriguingly, a significant increase in the proliferation of non-ventricular (or basal) progenitors was observed (Fig. 3F). The same analysis in *Chd8*^*neo/*-^ embryos revealed no significant differences in proliferation of either ventricular or non-ventricular progenitors (Fig. 3E-F), suggesting that the abnormal expansion of basal cortical progenitors only occurred within a restricted window of reduced CHD8 expression.

To determine the identity of the non-ventricular progenitors that proliferated more in mild hypomorphs, we stained serial sections with antibodies to SOX2, a marker of neural stem cells, TBR2, a marker for intermediate progenitors, and P3HB (Fig. 3G-J). The majority of proliferating cells were TBR2+ intermediate progenitors (70-80%), followed by TBR2+/SOX2+ cells (18-26%), most likely representing early intermediate progenitors, with a small minority (2-4%) of SOX2+/TBR2-cells, likely comprising the small population of stem-like, outer radial glia cells (oRGs) (Florio and Huttner, 2014; Hansen et al., 2010). There were no substantial changes in the proportions of mitotic, non-ventricular cell types between the different genotypes, indicating that a skewed distribution of non-ventricular zone progenitors was not responsible for the mitotic differences observed (Suppl. Fig. 3D-E).

Once again, increased numbers of mitotic, non-ventricular progenitors were observed in *Chd8^neo/neo^* embryos compared to wildtypes, but no difference between *Chd8*^*neo/*-^ embryos and their wildtype littermates (Fig. 3K). As expected, both TBR2-positive (Fig. 3L) and SOX2-positive cells (Fig. 3M) showed increased proliferation in *Chd8^neo/neo^* embryos. Consistent with our PH3B-only data (Fig. 3K), *Chd8*^*neo/*-^ embryos showed no changes in the number of proliferating TBR2-positive (Fig. 3L) or SOX2-positive (Fig. 3M) cells compared to wildtypes. In accordance with increased proliferation of TBR2+ progenitors, the number of TBR2+ cells was significantly increased in *Chd8^neo/neo^* embryos (Fig. 3N). Surprisingly, TBR2+ cell numbers were also slightly increased in *Chd8*^*neo/*-^ embryos, suggesting that these progenitors may be proliferating at slightly elevated levels in these mutants, although we could not accurately detect this difference using PH3B immunohistochemistry (Fig. 3K-L).

To understand the possible reason for the reduced cell proliferation in severe compared to mild hypomorphs, we interrogated gene expression data for genes or pathways that may underlie this difference. Comparing gene ontology analyses between these mice (Suppl. Fig. 4 and 5; Suppl. Table 3), we noticed a slight increase in the number of p53-regulated genes and ribosomal genes like *Rpl26* that can augment p53 mRNA translation (Takagi et al., 2005) in severe hypomorphs, raising the possibility that progenitors may be more prone to cell cycle exit and apoptosis in these mice. Indeed, cleaved caspase 3 (CC3) immunostaining revealed increased numbers of apoptotic cells within *Chd8*^*neo/*-^ embryos compared to *Chd8^neo/neo^* embryos (Fig. 3O-S), supporting this possibility.

### CHD8 expression is essential for repression of p53 target genes in early embryonic neocortex

Our analysis of *Chd8*^+/-^, *Chd8^neo/neo^* and *Chd8*^*neo/*-^ mice has revealed threshold transcriptional responses to decreasing CHD8 dosage, so that novel genes and pathways are dysregulated as CHD8 protein levels is reduced from ~50% to ~25%. To explore the consequences of complete CHD8 loss (0%), we conditionally deleted *Chd8* in neural progenitors. *Sox1*-*cre*-mediated deletion (Fig. 4A-B, Suppl. Fig. 6) of *loxP*-flanked (flox) exon 3 results in an early frameshift and termination of translation at amino acid 419, which is predicted to produce a protein that lacks all functional domains and results in a *Chd8*-null allele as shown previously in *Chd8*^+/-^ mice (Suetterlin et al., 2018).

**Fig 4.**
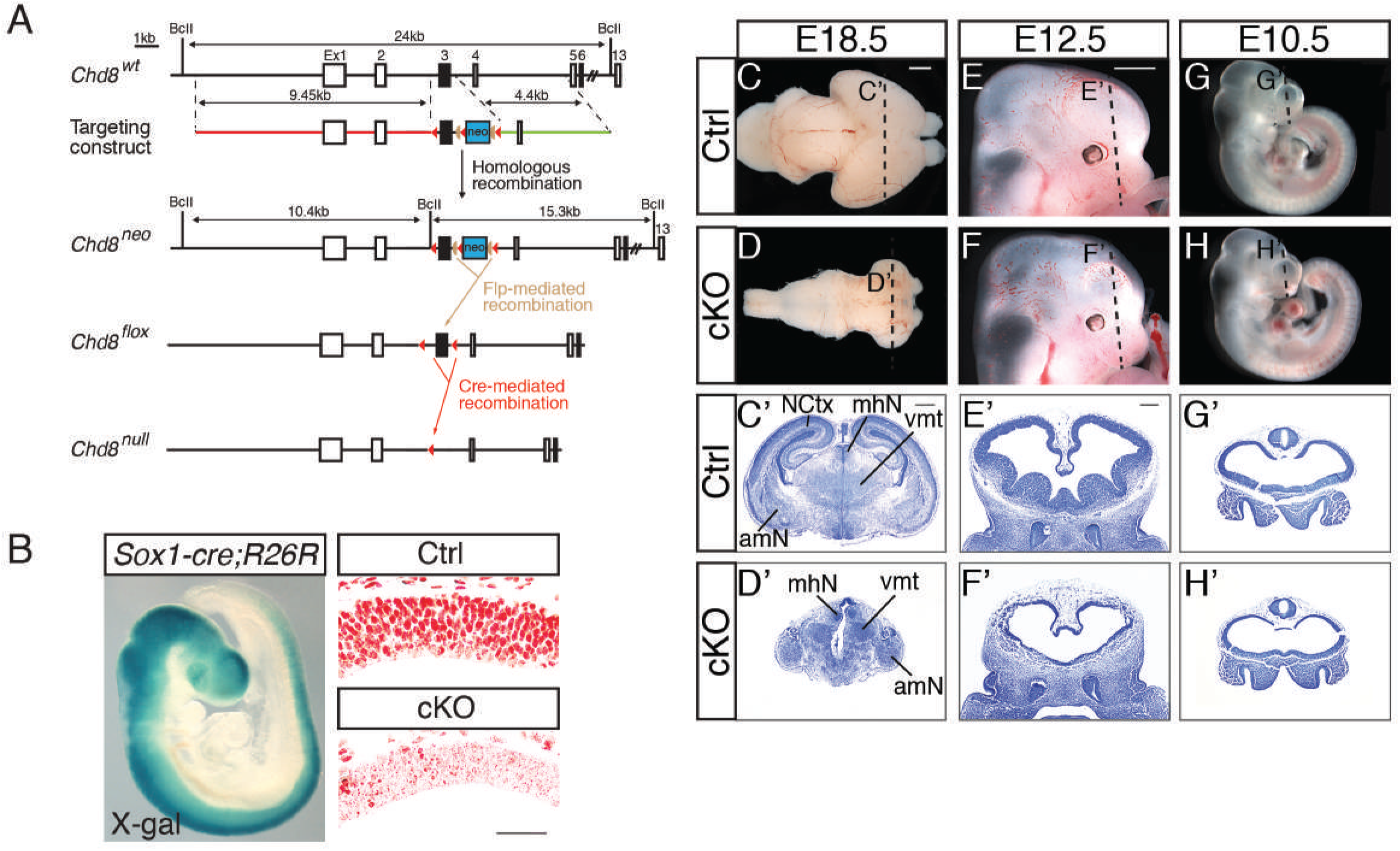
Conditional deletion of *Chd8* from the embryonic neuroepithelium results in severe hypoplasia of the telencephalon and neocortex. A)Schematic representation of the wildtype (wt) mouse *Chd8* gene (*Chd8^wt^*), targeting construct for homologous recombination in embryonic stem cells, the *Chd8* targeted allele (*Chd8^neo^*), the *Chd8* conditional allele after Flp-mediated excision of the neomycin resistance cassette (*Chd8^flox^*) and the *Chd8^null^* allele produced by Cre-mediated deletion of exon 3. Boxes represent exons, with exon 1 (Ex1) to 6 and 13 shown and exon 3 filled in black. The blue box represents a neomycin resistance cassette (neo), red triangles represent *loxP* sites and tan triangles *frt* sites. The long 9.45 kb (5’) homology arm is indicated in red and the short 4.4 kb (3’) homology arm in green in the targeting construct. X-gal staining of a *Sox1*-*Cre;R26R* embryo at E9.5 (left); and immunostaining for CHD8 protein on *Chd8^flox/flox^* (Ctrl) and conditional knockout *Sox1*-*Cre;Chd8^flox/flox^* (cKO) E10.5 neural tube (right). Scale bar = 50μm C,D) Wholemount images of E18.5 brains of a representative Ctrl and cKO embryo, anterior is to the right. Scale bar = 1mm E,F) Wholemount images of embryonic day 12.5 heads, anterior to the right. Scale bar = 1mm G,H) Wholemount images of E10.5 embryos, anterior to the right. C’ -H’) Cresyl violet-stained frontal sections through brains as indicated in C-H above. Scale bars = 500μm (C’-D’) and 200μm (E’-H’). The following subcortical structures are labelled in Ctrl (C’) and cKO (D’) at E18.5: NCtx: Neocortex, mhN: medial habenular nucleus, vmt: ventral medial thalamic nucleus, amN: amygdaloid nucleus.

The pan-neuronal, conditional deletion of *Chd8* by *Sox1*-*cre* (Fig. 4B, Suppl. Fig. 6) resulted in pronounced brain hypoplasia in homozygous conditional knockout (cKO) embryos, compared to controls (Ctrl, *Chd8^flox/flox^* (*Chd8^f/f^*)) (Fig. 4C-D). Neocortical hypoplasia, with the maintenance of some subcortical brain structures was evident upon histological analysis of E18.5 embryos (Fig. 4C’,D’). To identify the origin of these defects, cKO embryos were examined at earlier stages of development. Telencephalic hypoplasia with markedly thinner neuroepithelium was evident in E12.5 cKO embryos when compared to controls (Fig. 4E-F’). Examination of E10.5 cKO embryos showed telencephalic vesicles of near-normal size with neuroepithelia that were slightly thinner than controls (Fig. 4G-H’), suggesting that CHD8 is essential for expansion of the pallium from early embryonic development.

RNA-seq analysis identified 2032 DEGs in E10.5 cKO telencephalic vesicles compared to controls (Fig. 5A, Suppl. Table 4). KEGG pathway mapping of all dysregulated DEGs identified the p53 pathway as the most affected pathway (Fig. 5B, Suppl. Fig. 7). Interestingly, GO analysis identified cell cycle as the most dysregulated bioprocess (Suppl. Table 4), with a slight majority of genes within this category downregulated (60 out of 111). Quantitative RT-PCR (qRT-PCR) confirmed significant upregulation of multiple p53-regulated genes (Fig. 5C). Furthermore, genes normally upstream of p53, *Atr* and *Atm*, and *Trp53* (the gene encoding p53 itself) were not affected by *Chd8* deletion (Fig. 5C), consistent with a role for CHD8 in directly repressing p53 target genes (Nishiyama et al., 2009). To test whether p53 pathway hyperactivation was responsible for the cKO phenotype, we generated cKO embryos with reduced p53 gene dosage. Neocortical hypoplasia was partially rescued in *Sox1*-*Cre;Chd8^f/f^;Trp53*^*f/*+^ (conditional knockout p53 heterozygous, cKO-p53het) embryos (Fig.5D).

**Fig 5.**
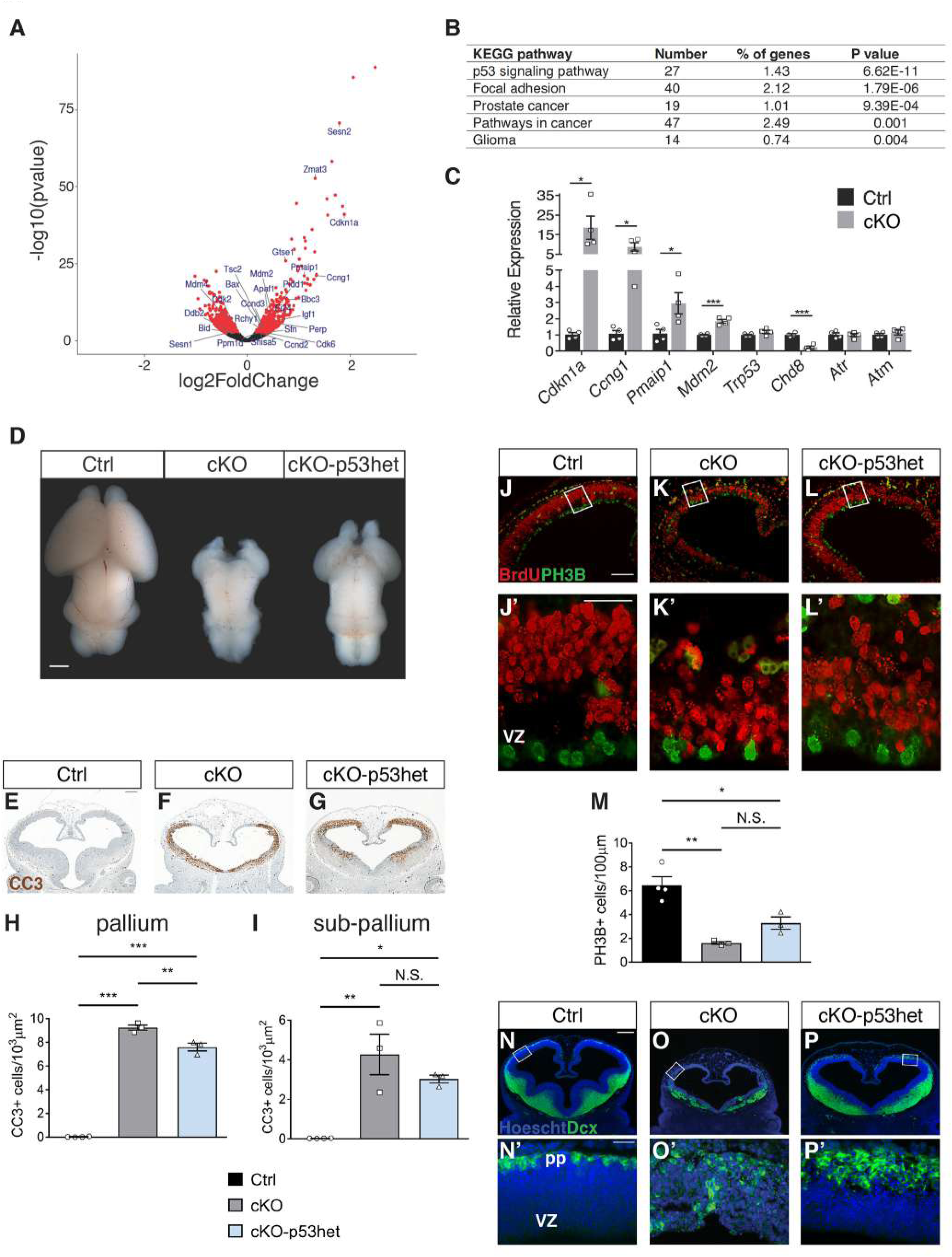
Repression of p53 target genes by CHD8 is necessary for normal brain growth. A)Volcano plot of RNA-seq data illustrating in red genes that are differentially expressed (FDR<0.05) in E10.5 cKO telencephalon, with p53 pathway genes labelled. Pathway enrichment analysis of differentially expressed genes, with the top KEGG pathway terms shown. See also Supplementary Table 3. qRT-PCR validation of a selection of p53 pathway genes identified by RNA-seq. (n=4 for each condition. Mean±SEM; *p<0.05, ***p<0.001, student’s t-test). D)Wholemount brains from E18.5 Ctrl, cKO and cKO-p53het mice are shown, anterior to the top. Data are representative of 6 embryos per genotype. Scale bar = 1mm E-G) Cleaved caspase 3 (CC3) immunohistochemistry (brown) on frontal sections through the telencephalon of E12.5 embryos. Scale bar = 200μm. H)Quantification of CC3-positive cells/μm^2^ of the pallium in embryos of each genotype (Ctrl, n=4; cKO, n=3; cKO-p53het, n=3; Mean±SEM; *p<0.05, **p<0.01, ANOVA followed by Tukey’s multiple comparisons test). I)Quantification of CC3-positive cells/μm^2^ of the subpallium (medial ganglionic eminence) in embryos of each genotype (Ctrl, n=4; cKO, n=3; cKO-p53het, n=3; Mean±SEM; *p<0.05,**p<0.01, ANOVA followed by Tukey’s multiple comparisons test). J-L) Immunohistochemistry to detect BrdU (red) and phospho-histone 3B (PH3B)-positive nuclei in frontal sections through the telencephalon of E12.5 embryos. Scale bar = 100μm. J’-L’) Magnified images of the boxed neocortical regions in J-L, with the ventricular zone (vz) at the bottom and pial surface at the top. Scale bar = 25μm (M) Quantification of PH3B-positive cells/μm of neocortical ventricular surface (Ctrl, n=4; cKO, n=3; cKO-p53het, n=3; Mean±SEM; *p<0.05, **p<0.01, ANOVA followed by Tukey’s multiple comparisons test). N-P) Immunostaining of frontal E12.5 sections for DCX (green) to label differentiating neurons, with nuclei counterstained with Hoechst 33342. Scale bar = 200μm. N’-P’) Magnified images of boxes in N-P, with the ventricular zone (vz) at the bottom and pial surface (pp) at the top. Scale bar = 25μm.

A substantial increase in apoptosis was observed in the cKO embryos (Fig. 5E,F,H,I). Although the number of apoptotic cells seen in the cKO-p53het was significantly reduced in the pallium compared to the cKO (Fig. 5F-I), the level of cell death was still considerably greater than that seen in controls, indicating that p53 heterozygosity was not sufficient to rescue the apoptotic phenotype in cKO embryos.

We also noted the presence of certain cell cycle inhibitors amongst genes upregulated in the cKO (e.g. p21/CDKN1A)(Fig. 5C). Therefore, we investigated neural progenitor proliferation. BrdU incorporation was used to identify neural progenitors in S phase, and PH3B labelling to visualise mitotic progenitors. Fewer BrdU-positive cells were present in cKO embryos and quantification of PH3B-positive cells in the ventricular zone of the neocortex confirmed a strong reduction in cell proliferation (Fig. 5J,J’,K,K’,M). Progenitor proliferation was not significantly rescued in cKO-p53het embryos compared to cKO embryos (Fig. 5L,L’,M), consistent with only a partial rescue of neocortical size in these animals (Fig. 5D).

The increase in cell death and reduced proliferation of progenitors in cKO neocortex suggested that many of these progenitors were prematurely exiting the cell cycle. To examine how this might affect the production of postmitotic neurons, we visualised neuronal differentiation by immunostaining for Doublecortin (DCX). DCX+ cells accumulated at the pial surface in control embryos (Fig. 5N). The first DCX+ differentiating neurons of the pallium were clearly visible in the preplate (pp) (Fig. 5N’). Ectopic clusters of DCX+ cells were visible throughout the cKO neural tube, including the ventricular zone (Fig. 5O,O’), suggesting that some progenitors were precociously differentiating. Whereas DCX+ cell positioning was normalised in cKO-p53het embryos, the enhanced differentiation of cells was not fully rescued, with DCX+ cells encompassing almost 50% of the pallium (Fig. 5P,P’).

Taken together, these data identify CHD8 as an essential repressor of p53 pathway activation during neocortical development. CHD8 loss leads to increased apoptosis, reduced neural progenitor proliferation and precocious cell differentiation during early embryonic development, resulting in severe neocortical hypoplasia by the end of gestation.

## Discussion

Human genetic studies have identified heterozygous, likely gene disrupting mutations in *CHD8* as a possible cause of ASD and macrocephaly. *Chd8* heterozygous mice have been generated by several groups, but these mice were found to exhibit relatively subtle brain overgrowth (Gompers et al., 2017; Jung et al., 2018; Katayama et al., 2016; Platt et al., 2017). Observations of relatively small transcriptional changes in the mid-gestation *Chd8*^+/-^ mouse brain appeared at odds with the many genes dysregulated upon *Chd8* knockdown in progenitor cell lines and after in utero electroporation (Cotney et al., 2015; Durak et al., 2016; Sugathan et al., 2014).

Together, these studies led us to explore whether different sensitivities to reduced CHD8 dosage might account for some of these inconsistencies. A comparison of brain size, gene expression and neural progenitor fate in a mouse *Chd8* allelic series yielded several key findings: 1) A small additional reduction in *Chd8* expression in mild hypomorphs compromised the capacity of neural progenitor cells to maintain stable expression of over 2200 genes in the mid-embryonic neocortex, which included over 150 ASD-associated genes. *Chd8^neo/neo^* mice also exhibited more pronounced brain hyperplasia (Fig. 6). 2) We identified a key role for CHD8 in limiting the expansion of TBR2+ intermediate progenitors, a population particularly important for human cortical development. 3) We report several examples of reduced CHD8 expression having non-linear effects (Fig. 6). In addition to the precipitous gene expression changes in mild hypomorphs, we observed a striking activation of p53-regulated genes, precocious neural progenitor differentiation and abnormal levels of apoptosis resulting in severe brain hypoplasia upon homozygous *Chd8* deletion in neural progenitors. Together, these findings indicate that CHD8 levels need to be tightly regulated during development and that the interpretation of experimental manipulations that involve *Chd8* knock-down should consider these non-linear, threshold effects. It seems likely that the reduced neural progenitor proliferation reported upon *Chd8* knockdown by Durak et al. is a result of reducing CHD8 protein levels by 80%, rather than 50% (Durak et al., 2016). The possibility that some human mutations in ASD patients may result in dominant negative proteins leading to more pronounced phenotypes needs to be explored.

**Fig 6.**
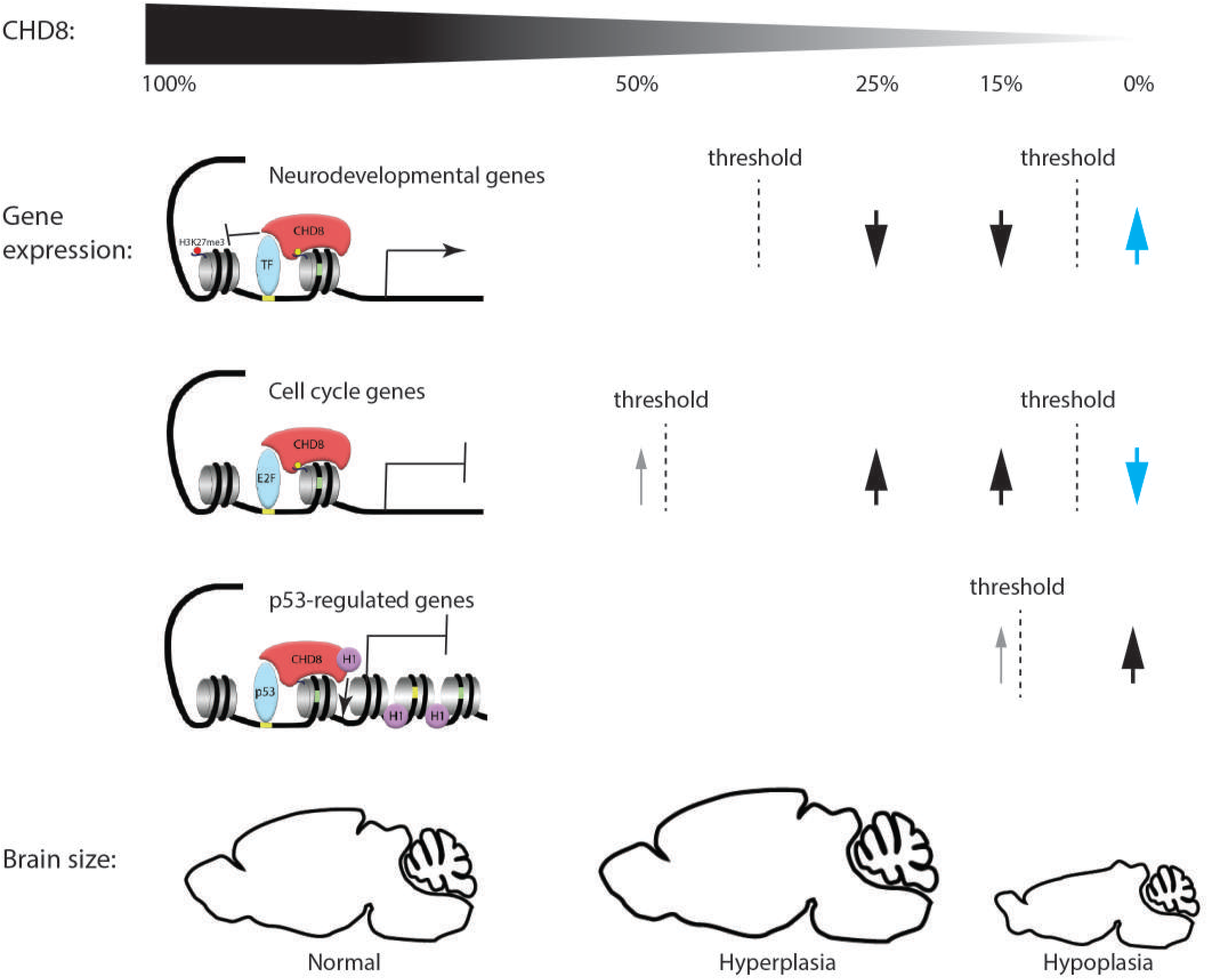
The non-monotonic relationship between CHD8 protein levels, gene expression and brain size. The effects of stepwise reductions in CHD8 protein levels to ~50% (heterozygous), ~25% (mild hypomorph), ~15% (severe hypomorph) and 0% (conditional knockout) on the transcription of neurodevelopmental, cell cycle and p53-regulated genes and brain size are depicted. CHD8 appears to function primarily as a positive regulator of neurodevelopmental genes via recruitment to H3K4me3-modified (yellow ball) histones (gray spool), presumably via enabling the recruitment of key transcription factors (TF) and antagonising Polycomb-mediated repression (H3K27me3, red ball). A sharp reduction in the expression of many of these genes (arrow) is only observed in E12.5 neocortex when CHD8 levels are reduced to below a threshold less than haploinsufficient levels. CHD8 appears to repress E2F-regulated cell cycle genes in this context, with significant induction only becoming evident at sub-haploinsufficient levels, although low expression increases (grey arrow) likely drives subtle increases in proliferation in the heterozygous state. Both neurodevelopmental and cell cycle genes are dysregulated in the opposite direction in the cKO, suggestive of non-monotonic effects (blue arrows). CHD8 can interact with p53 and histone 1 (H1), leading to stable heterochromatin formation and repression of p53 target genes. A few p53-regulated genes become activated in hypomorphic mice (grey arrow), but the majority remains fully repressed with de-repression only becoming evident upon complete CHD8 loss. Note the different CHD8 thresholds for different groups of genes (broken lines) and the non-monotonic effects on gene expression and over-all brain size.

### CHD8 as a phenotypic capacitor

It has been posited previously that ASD-associated chromatin remodelling factors may act as phenotypic capacitors, buffering against perturbations to normal development in order to maintain stable phenotypes (Suliman et al., 2014). Heterozygosity for a capacitor is predicted to result in a loss of robustness, such that the individual is more susceptible to additional genetic and non-genetic risk factors. Our findings that over 2200 genes, many of which are ASD risk factors, became dysregulated by a small additional decrease in CHD8 dosage below 50%, supports the idea that many neurodevelopmental genes and processes may be inherently unstable in the *Chd8* heterozygous neocortex.

This hypothesis also suggests the possibility that brain development in human and mouse differ in robustness, such that *CHD8* heterozygosity may cause more pronounced changes to brain growth and transcriptional regulation in the developing human brain. However, the actual effects of reported ASD-associated *CHD8* mutations on CHD8 protein levels and CHD8 function remain to be determined. The possibility that these mutations reduce CHD8 protein levels to less than 50% or lead to dominant negative protein remains. Conversely, the C57BL/6 genetic background used for all *Chd8* heterozygous mouse studies so far, may be protective and more robust, and different phenotypes may emerge on different genetic backgrounds, as might be expected for a phenotypic capacitator.

### CHD8 regulates the proliferation of non-ventricular cortical progenitors

Our observation of increased proliferation of neural progenitors outside the ventricular zone in the embryonic neocortex of *Chd8^neo/neo^* mice suggest the possibility that fundamental differences in mouse and human brain development may result in *CHD8* haploinsufficiency having more pronounced effects on human brain development. Comparative studies of gyrencephalic and lissencephalic animals have identified important differences in the capacity of non-ventricular progenitors to expand and subsequently contribute to cortical expansion. This population of progenitors consists of outer radial glia cells (oRGs), which express *Sox2* and typically retain a characteristic basal process, and *Tbr2*-expressing intermediate progenitor cells (IPs) (Florio and Huttner, 2014). In humans, oRGs are located in an expanded outer-subventricular zone (oSVZ) and are capable of asymmetric divisions that generate an oRG daughter cell, which maintains the pool of non-ventricular progenitors, and an IP daughter cell that can undergo transit-amplifying divisions to expand and generate neuronal progeny (Hansen et al., 2010). By contrast, mouse oRGs primarily undergo self-renewing, neurogenic divisions and populate a non-ventricular region lacking the distinct, expanded cytoarchitecture of the oSVZ typically seen in gyrencephalic species (Wang et al., 2011). Furthermore, mouse IPs likely possess a more limited capacity for self-renewal, as a majority of mouse IP divisions appear to generate two neuronal daughter cells (Kowalczyk et al., 2009; Miyata et al., 2004). Therefore, if CHD8 has an especially crucial role in regulating the expansion of this specific population of neural progenitors, then it is possible that CHD8 deficiency in these cells could result in more pronounced phenotypes with regard to cortical over-growth and circuit disruption in humans. Interestingly, we also note that Bernier et al. previously identified an enrichment for CHD8 expression in areas outside the ventricular zone in human mid-fetal cortex (Bernier et al., 2014), further supporting the idea that CHD8 may have an important role in regulating expansion of these cells.

### CHD8 is an essential repressor of p53 in neural progenitors

One of the most striking non-monotonic effects of *Chd8* efficiency reported here is brain hypoplasia identified in pan-neuronal *Chd8* cKO mice, partly as a result of de-repression of the p53 pathway. This discovery identifies CHD8 as a critical repressor of p53 target gene activation in neural progenitors. Our findings suggest that very low levels of CHD8 are sufficient to repress p53 target genes and maintain neural progenitor self-renewal. One could speculate that the CHD8-dependent recruitment of histone H1 to p53 target genes to initiate a cooperative process of chromatin compaction (Nishiyama et al., 2009), may require lower levels of CHD8 than another process that is dependent upon the constitutive recruitment of RNA polymerase (Rodriguez-Paredes et al., 2009) or other co-activating factors by CHD8 (Fig. 6). Our gene expression and apoptosis data suggest that CHD8 protein levels in *Chd8*^*neo/*-^ embryos were close to this critical threshold. Cotney et al. reported p53 signaling as one the most dysregulated pathways upon CHD8 knock-down in human neural stem cells (Cotney et al., 2015). However, other studies in human cell lines have not demonstrated the same changes, including an in vitro knock-down of CHD8 to 20-25% of control levels in human SK-N-SH neural progenitor cells (Wilkinson et al., 2015). Together, these findings suggest that transcriptional responses to reduced CHD8 levels are highly context-dependent and may help shed light on reports that *Chd8* knock-down in utero led to reduced proliferation and enhanced differentiation of neural progenitor (Durak et al., 2016). The knock-down of CHD8 protein levels to 20% of wildtype levels in this study may explain the discrepancies with the findings reported here and in other studies (Gompers et al., 2017; Katayama et al., 2016; Platt et al., 2017).

In conclusion, our analysis of an allelic series of *Chd8*-deficient mice has identified clear non-monotonic effects on gene expression and brain development (Fig. 6). Recognition of the differing sensitivities of important cellular processes to CHD8 dosage is an important step in understanding the context-specific transcriptional roles of CHD8 and explain how relatively small differences in CHD8 levels may lead to disproportionally large differences in phenotype.

## Experimental Procedures

### Mice

A transgenic mouse line containing a *Chd8^neo^* allele (*Chd8^tm1.Mabn^*) was generated as reported previously (Suetterlin et al., 2018). Briefly, an 18.8 kb targeting construct was generated consisting of a 14.84kb genomic DNA fragment subcloned from a C57BL/6 BAC clone (RP23:318M20) with an added loxP/FRT-PGK-gb2-Neo cassette 3’ of exon 3 (ingenious Targeting Laboratory (iTL), Ronkonkoma, NY, USA) and additional loxP site 5’ of exon 3 (Fig. 1). The targeting construct was linearised and electroporated in C57BL/6J ES cells. Five clones were identified with successful recombination, two of which (124 and 254) were injected into Balb/c blastocysts. Resulting chimaeras were backcrossed onto a C57BL/6J background to generate *Chd8*^*neo/*+^ mice. Experimental *Chd8^neo/neo^* mice were produced by *Chd8*^*neo/*+^ × *Chd8*^*neo/*+^ crosses. To generate a conditional *Chd8* allele (*Chd8^flox^* (*Chd8^tm1.1Mabn^*)), *Chd8*^*neo/*+^ mice were crossed with Flpe deleter mice on a C57BL/6J background (Fig. 1). *Chd8*^*flox/*+^ mice were then either inter-crossed to obtain a homozygous *Chd8^flox/flox^* line or with *Sox1*-*Cre* (Arnold et al., 2008) to generate *Sox1*-*Cre; Chd8*^*flox/*+^ mice. To produce pan-neuronal *Chd8* null (conditional knockout, cKO) mice, *Sox1*-*Cre; Chd8*^*flox/*+^ mice were mated with *Chd8^flox/flox^* mice. *Sox1*-*Cre; Chd8^flox/flox^* cKO embryos were compared with *Sox1*-*Cre; Chd8*^*flox/*+^ (cHET) and *Chd8^flox/flox^* (Ctrl) embryos. To generate conditional p53-heterozygotes, mice carrying the *Trp53^tm1Brn^* conditional (*p53^flox^*) allele were obtained from the Jackson laboratories (Marino et al., 2000) and crossed to the *Chd8* conditional mice. *Chd8*^*flox/*+^ mice were also bred with *β*-*actinCre* mice (Lewandoski and Martin, 1997) to generate a *Chd8* null (*Chd8*^-^, (*Chd8^tm1.2Mabn^*)) allele. *β*-*actinCre;Chd8*^+/-^ mice were then crossed with C57BL/6J mice to remove the Cre transgene and establish a *Chd8*^+/-^ line. *Chd8*^+/-^ mice were produced by *Chd8*^+/-^ × C57BL/6J crosses, taking care to equalise paternal or maternal inheritance of the *Chd8* null allele. All animal procedures were approved by the UK Home Office.

Genomic DNA was extracted for genotyping from ear samples (or yolk sac for embryos aged E14.5 and below) using Proteinase K digestion or the HotSHOT method (Truett et al., 2000). Genotyping reactions were then performed for the presence of *Chd8* wildtype, null or floxed alleles, p53 wildtype or floxed alleles, as well as the presence of Cre. Thermal cycles for all genotyping reactions were as follows: 94°C, 5 minutes; 35X (94°C, 30sec; 58°C, 30sec; 72°C, 30sec); 72°C, 5 minutes. Primer pairs to amplify a sequence distinguishing between *Chd8^flox^*,*Chd8^neo^* and wildtype alleles (‘*Chd8flox*’ primers, 212bp and 275bp product for mutant or wildtype, respectively), to detect the presence of the *Chd8^null^* allele (‘*Chd8null*’ primers, 395bp), to distinguish between p53 floxed and wildtype alleles (‘*p53flox*’ primers, 390bps and 270bps, respectively) and primers to amplify a specific Cre sequence (‘*Cre*’ primers, 390bp product; see also figure S1) were used as indicated below:

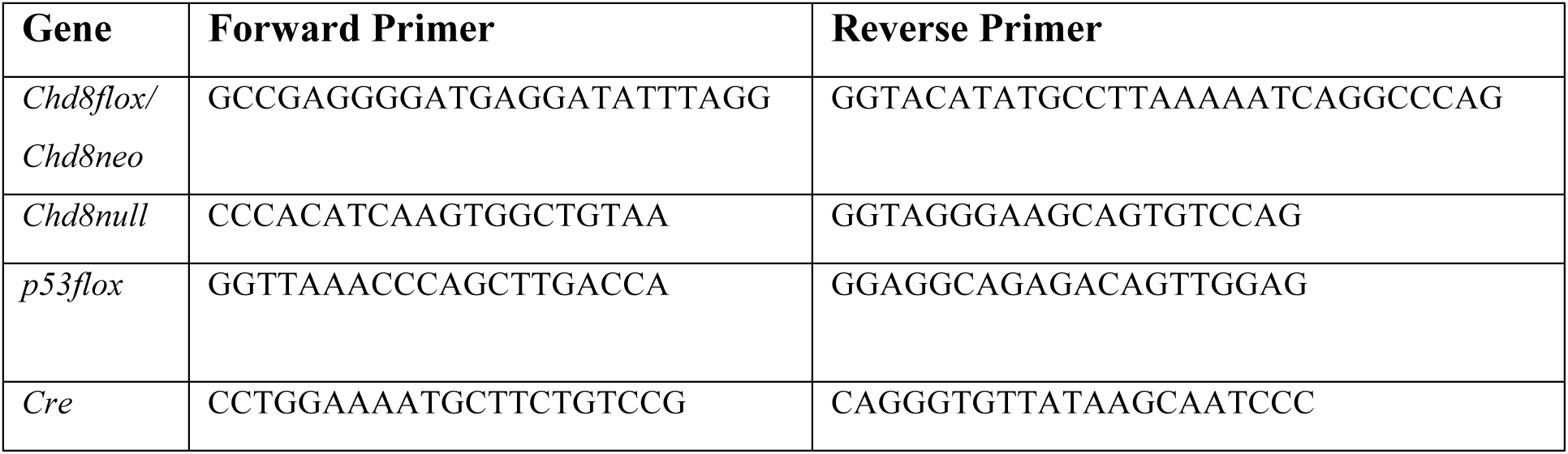

### RNA extraction and qRT-PCR analysis

Cortical RNA was extracted by lysing cortices in 600μl Trizol (Life Technologies). After purification, RNA was DNase treated using the Direct-zol RNA MiniPrep kit (Zymo Research) according to the manufacturer’s recommendations. cDNA was synthesised for qRT-PCR experiments using 50ng RNA from 4 biological replicates per condition with the Precision nanoScript 2 Reverse Transcription Kit (PrimerDesign Ltd.) according to the manufacturer’s instructions. qRT-PCRs were performed on a Stratagene Mx3000p (Agilent Technologies) using PrecisionPlus-MX 2x qPCR Mastermix with SYBR green (PrimerDesign Ltd.) and primers against *Atm*, *Atr*, *Trp53*, *Cdkn1a*, *Ccng1*, *Mdm2*, *Chd8*, *and Pmaip1*. Relative expression levels were calculated using the 2^-ΔΔCT^ method and *Gapdh* and *Ywhaz* were used as endogenous control genes.

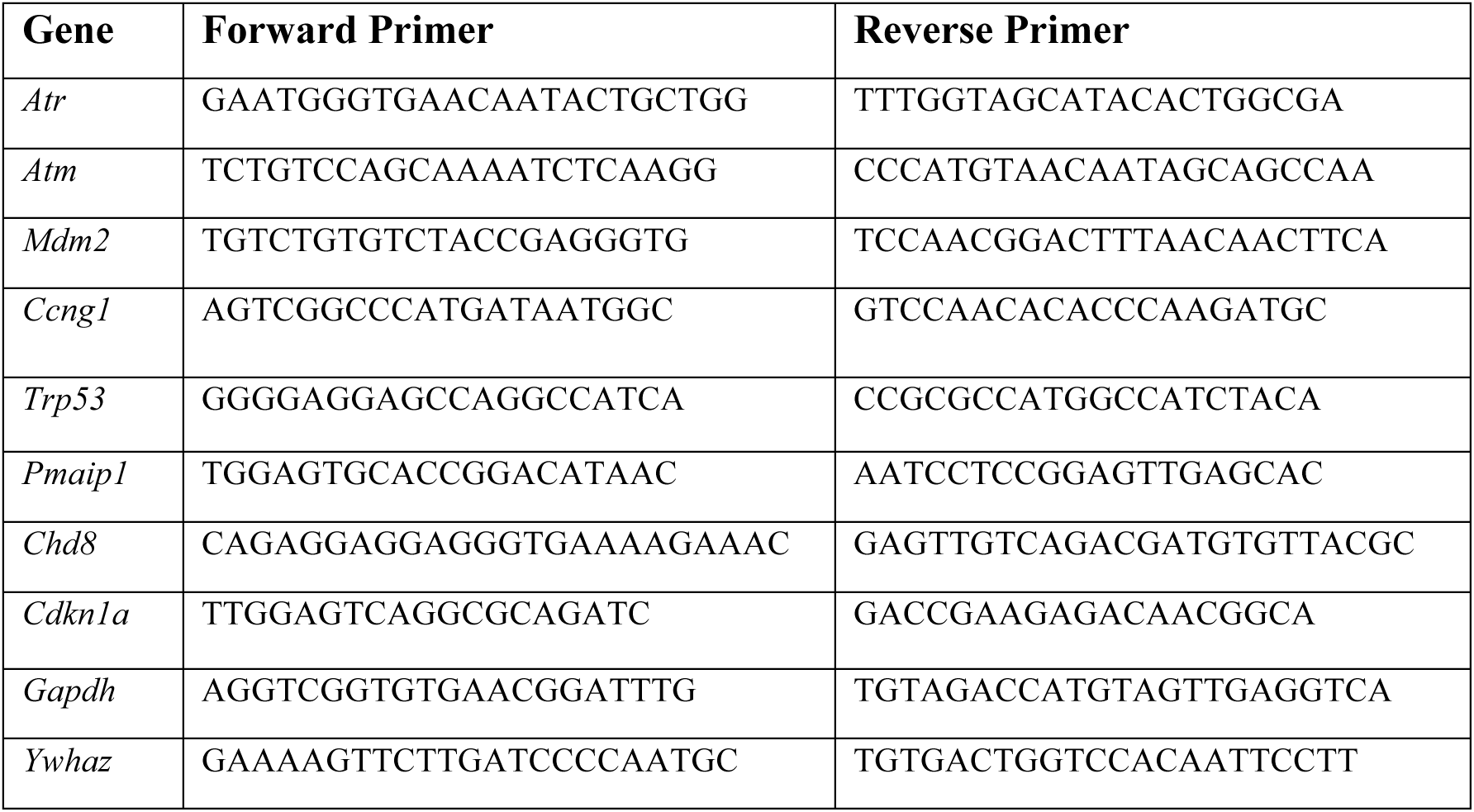
qRT-PCR primer sequences:

### Western blot

Telencephalic vesicles were dissected from E12.5 embryos and whole cell protein prepared by lysing in 8M urea, 1% CHAPS, 50mM Tris (pH 7.9) lysis buffer containing protease inhibitors (PMSF, Pepstatin A, Leupeptin, Aprotinin; Roche) and a phosphatase inhibitor cocktail (Sigma). After rotating at 4°C for 30 mins, DNA was removed from lysates by centrifugation. Supernatant was transferred to a fresh tube and stored at −80°C. Protein loading samples were made by diluting samples in Laemmli buffer containing 10% β-mercaptoethanol, followed by boiling at 100°C for 10 minutes. Samples were loaded (10μg total protein per lane) onto a Mini-PROTEAN pre-cast gel (Bio-Rad) and resolved using gel electrophoresis. Protein was transferred to a nitrocellulose membrane (Bio-Rad), which was then blocked in 5% non-fat milk powder (Bio-Rad) and 1% bovine serum albumin (BSA, Sigma) in TBS with 0.1% Tween-20 (TBST) for one hour at room temperature. Where β-actin was used as a loading control, the membrane was then cut in two: the higher molecular weight section was incubated with anti-CHD8 primary antibody (rabbit anti-CHD8 N-terminal, Bethyl Laboratories, 1/2000) and the lower molecular weight section incubated with anti-β-actin antibody (rabbit anti-β-actin, Abcam, 1/4000); both antibodies in 3% non-fat milk powder and 1% BSA in TBST overnight at 4°C. After washing, membrane was incubated with HRP-conjugated secondary antibody (Millipore) for one hour at room temperature. HRP was detected with Clarity ECL reagent (Bio-Rad) and the membrane imaged using a Bio-Rad ChemiDoc system. Where GAPDH was used as a loading control, the uncut membrane was washed in TBST after detection of CHD8 protein and incubated overnight at 4°C in 0.05% sodium azide in PBS, before washing and incubation with anti-GAPDH primary antibody (rabbit anti-GAPDH, Abcam, 1/40000) overnight at 4°C. Membrane was probed with HRP-conjugate and imaged as before. Relative protein quantity was calculated using Bio-Rad ImageLab software.

### Structural MRI

Mice were terminally anesthetized and intracardially perfused with 30mL of 0.1M PBS containing 10U/mL heparin (Sigma) and 2mM ProHance (a Gadolinium contrast agent) followed by 30mL of 4% paraformaldehyde (PFA) containing 2mM ProHance (Spring et al., 2007). Perfusions were performed at a rate of approximately 60mL/hr. After perfusion, mice were decapitated. The brain and remaining skull structures were incubated in 4% PFA + 2mM ProHance overnight at 4°C then transferred to 0.1M PBS containing 2mM ProHance and 0.02% sodium azide for at least 7 days prior to MRI scanning. A multi-channel 7.0 Tesla MRI scanner (Varian Inc., Palo Alto, CA) was used to image the brains within skulls. Sixteen custom-built solenoid coils were used to image the brains in parallel (Bock et al., 2005). Parameters used in the anatomical MRI scans: T2-weighted 3D fast spin-echo sequence, with a cylindrical acqusition of k-space, and with a TR of 350 ms, and TEs of 12 ms per echo for 6 echoes, two averages, field-of-view of 20 × 20 × 25 mm^3^ and matrix size = 504 × 504 × 630 giving an image with 0.040 mm isotropic voxels (Lerch et al., 2011). The current scan time required for this sequence is ~14 hours. To visualise and compare any changes in the mouse brains the images were linearly (6 parameter followed by a 12 parameter) and non-linearly registered towards a pre-existing atlas (Dorr et al., 2008), and then iteratively linearly and non-linearly aligned to each other to create a population atlas representing the average anatomy of the study sample. The result of the registration is to have all scans deformed into alignment with each other in an unbiased fashion. This allows for the analysis of the deformations needed to take each individual mouse’s anatomy into this final atlas space, the goal being to model how the deformation fields relate to genotype (Lerch et al., 2008; Nieman et al., 2006). The jacobian determinants of the deformation fields were then calculated as measures of volume at each voxel. Significant volume changes were then calculated in two ways, 1) on a region basis, and 2) voxelwise. Regional volumes are calculated by warping a pre-existing classified MRI atlas onto the population atlas. This atlas encompasses 159 different structures including, but not limited to, the cortical lobes, large white matter structures (i.e. corpus callosum), ventricles, cerebellum, brain stem, and olfactory bulbs (Dorr et al., 2008; Steadman et al., 2014; Ullmann et al., 2013). Significant differences can then be determined between groups for the 159 different regions in the brain. Voxelwise comparisons were made between mutants and all wildtypes taken from both the *Chd8*^+/-^ and *Chd8^neo/neo^* batches. As wildtype brains differed significantly between the two groups, most likely as a result of slightly different age at analysis, *Chd8^neo/neo^* data was first normalised (beta-corrected) to wildtypes in the *Chd8*^+/-^ batch before analysis. Voxelwise comparisons were then made between mutants and all wildtypes, and multiple comparisons in this study were controlled for using the False Discovery Rate (Genovese et al., 2002).

### RNA Sequencing

For RNA-sequencing at E10.5, total RNA from 2 animals was pooled for each biological replicate (n=3 per condition). No pooling was performed at E12.5 (n=3 per condition). mRNA was isolated and reverse transcribed into cDNA. cDNA was end-repaired, adaptor-ligated and A-tailed. Paired-end sequencing was performed on the Illumina HiSeq 4000 platform. Quality of the raw sequencing data was checked using FastQC version 0.11.2 and trimming of adaptor sequences was performed using Trim Galore! version 0.4.1 (Krueger et al., 2012). Reads were aligned to the mouse genome (GRCm38.p4) using Tophat version 2.1.0 and aligned reads were counted using FeatureCounts version 1.5.0 (Kim et al., 2013; Liao et al., 2014). Differential expression testing was performed using DESeq2 version 1.10.1, as previously described (Love et al., 2014). Gene ontology analysis and functional classification was performed using DAVID with all detected DEGs below a 0.05FDR (Huang da et al., 2009). For heatmaps, data were transformed with a variance stabilising transformation, scaled and clustered with the Ward.d2 method using maximum distance, and plotted with the R package pheatmap version 1.0.8. The R package ggplot2 version 2.1.0 was used to generate volcano plots and DESeq2 was used to generate normalised read count plots for individual genes. The list of ASD associated genes used for overlap with the neo/neo DEGs was obtained from the SFARI Human Gene database (https://gene.sfari.org/autdb/HG_Home.do). RNA-seq data have been deposited into GEO, accession number GSE81103.

### Tissue collection and processing

Embryos were collected by dissection, followed by dissection of brains from the skulls in ice-cold PBS for E18.5 embryos. Wholemount pictures were taken on a Nikon SMZ1500 stereomicroscope equipped with a Nikon DS-Fi1 camera head, followed by post-fixation in 4% PFA for 24h at 4°C. For BrdU experiments, pregnant mothers were injected with 40mg/kg BrdU in 0.9% saline 1 hour (for cKO proliferation assays) or 20 hours (for *Chd8^neo/neo^* proliferation assays) prior to death. After fixing, embryos were dehydrated and paraffin embedded. Paraffin blocks were then cut into 10μm (cKO embryos) or 5μm (*Chd8^neo/neo^* embryos) thick coronal sections and mounted such that each slide contained three adjacent sections.

#### X-gal staining

E9.5 embryos were collected and dissected in ice-cold PBS and post-fixed in 4% PFA for 10 minutes. Following three washes in PBS (5 minutes each), embryos were incubated in X-Gal staining solution (10mM TRIS-HCL, pH7.3, 0.005% Na-deoxycholate, 0.01% IGEPAL, 5mM potassium ferrocyanide, 5mM potassium ferricyanide, 2mM MgCl_2_, 0.8mg/ml X-Gal, in PBS) at room temperature until adequate signal was observed. Reactions were stopped by washing in PBS (3 × 5 minutes) followed by post-fixation in 4% PFA for 1h. Control embryos never showed any staining.

#### Immunohistochemistry and Immunofluorescence

Coronal brain sections were re-hydrated using standard protocols. Antigen retrieval was conducted by heating slides in 10mM Sodium Citrate solution (pH6) for 20mins and cooled on ice. For non-fluorescence immunohistochemistry, endogenous peroxidases were blocked by incubating in 3% H_2_O_2_ and 10% MeOH in PBS for 15mins. Sections were then washed in 0.2% Triton X-100 (Sigma-Aldrich) in PBS (PBT2) for 5 mins and blocked using 10% heat-inactivated normal goat serum (GS) and 2% gelatin in PBT2 for 1 hour. Sections were incubated in 5% GS in PBT2 containing primary antibody overnight at 4°C. The following antibodies were used: mouse anti-BrdU (BD Biosciences, 1/100), rabbit anti-phosphohistone 3B (Cell Signaling, 1/100), mouse anti-phosphohistone 3B (Abcam, 1/200), rabbit anti-sox2 (Abcam, 1/100), chicken anti-Tbr2 (Merck Millipore, 1/200), rabbit anti-Ki67 (Abcam, 1/200), rabbit anti-cleaved-caspase 3 (Cell Signaling, 1/200), rabbit anti-doublecortin (Abcam, 1/400) or rabbit anti-CHD8 (Bethyl, 225A, 1/400). For immunofluorescence, sections were incubated with secondary antibody diluted in 5% GS in PBT2 for 90mins at 4°C. Secondary antibodies used included goat anti-chicken AlexaFluor 488 (Invitrogen, 1/200), goat anti-mouse AlexaFluor 405 (Invitrogen, 1/200), goat anti-mouse AlexaFluor 594 (Invitrogen, 1/200), goat anti-rabbit AlexaFluor 488 (Invitrogen 1/200), goat anti-rabbit AlexFluor 658 (Invitrogen, 1/200) and donkey anti-rabbit AlexaFluor 488 (Invitrogen, 1/200). Sections were then counterstained using Hoechst 33342 solution (Invitrogen, 1/50,000) in PBS and then mounted onto coverslips using CitiFluor (CitiFluor Ltd., UK). For diaminobenzidine (DAB) immunohistochemistry, after incubation with primary antibody sections were incubated in biotinylated anti-rabbit immunoglobulin secondary antibody (Dako, 1/200) in 5% GS in PBT2. Samples were washed in PBS and incubated with Avidin/biotin complex (ABC, Vector) in PBS for 1 hour. Sections were developed using 0.03% DAB and 0.0003% H_2_O_2_ in PBS for 10mins before washing in running water and counterstaining using Ehrlich’s Haemotoxylin solution. Slides were mounted onto coverslips using DPX (Sigma-Aldrich). Images were acquired on a Nikon 80i microscope equipped with a Hamamatsu C4742 CCD or Nikon 5M pixel Nikon DS digital cameras. Images were processed using Adobe Photoshop and Illustrator.

#### Quantitative Analysis

Proliferation was quantified by counting phosphohistone 3B-positive cells either lining the ventricular surface of the dorsal cortex (‘ventricular’) or basal to the ventricular zone (‘non-ventricular’) and these counts normalised to the length of ventricular surface. These were quantified and averaged across both hemispheres in at least three consecutive sections and averaged to calculate the number of phosphohistone 3B-positive cells per μm of ventricular surface in the dorsal cortex. Tbr2-positive, Sox2-positive and double-positive cells within the subventricular were counted within 150μm × 40μm boxes placed over the subventricular zone, which was determined based on Tbr2-positivity. This was performed in three separate locations within each hemisphere: i) a ‘dorsal’ region located at the apex of the dorsal neocortex, ii) an ‘intermediate’ region located at approximately 50% of neocortex length and iii) a ‘ventral’ region located just dorsal to where the neocortex meets the lateral ganglionic eminences. Counts for each separate region were averaged across both hemispheres in at least three consecutive sections per biological replicate. As counts in each individual region showed no significant difference from each other (see S4 Fig), these were summed together for each biological replicate to give total counts per 1800μm^2^ of subventricular zone. For Q fraction experiments, 200μm segments of neocortex were measured using ImageJ. In these segments, BrdU-positive/Ki67-negative cells were first counted basal to the subventricular zone, followed by quantification of all BrdU-positive cells within the segment. The fraction of progenitors that quit the cell cycle (Q fraction) was then defined as the proportion of BrdU-positive/Ki67-negative cells accounting for all BrdU-positive cells within the segment. For each biological replicate the Q fraction was averaged across both hemispheres of a given section, followed by averaging across at least three consecutive sections. Apoptosis in dorsal cortex was quantified by counting cleaved-caspase 3 (CC3)-positive cells in 50μm × 50μm boxes. Three boxes were counted for both inner (ventricular side) and outer (pial side) regions of the dorsal cortex to generate an average number of CC3-positive cells per μm^2^ for both inner and outer cortical regions, which were then averaged to provide the overall mean of CC3-positive cells per μm^2^. These were calculated for both sides of the brain individually in at least two adjacent sections per biological replicate. Apoptosis in the ventral cortex was quantified by counting CC3+ cells either in three 0.1mm × 0.1mm boxes in both lateral ganglionic eminences and three 0.15mm × 0.15mm boxes in both medial ganglionic eminences (WT and cHet), or three 0.1mm × 0.1mm boxes placed at equivalent positions across the ventral cortex (cKO and cKO-p53Het) and counts averaged for each section. These were calculated for both sides of the brain individually in at least two adjacent sections per biological replicate. All data were analysed using GraphPad Prism 6 and significance calculated using student’s t-test.

## Author contributions

SH, CM, PS, JE and FR designed and performed experiments, analysed the data and produced figures. JPL, CF and MAB supervised the experimental work and analyses. SH, CM, PS and MAB wrote the manuscript with input from all authors. The study was conceived by MAB.

## Acknowledgements

This work was supported by research grants from the Medical Research Council (MR/K022377/1, MAB and CF), Simons Foundation (SFARI #344763, MAB) and Ontario Brain Institute’s POND programme (JPL). SH was supported by the King’s Bioscience Institute and the Guy’s and St Thomas’ Charity Prize PhD Programme in Biomedical and Translational Science. We thank Elizabeth Robertson (University of Oxford) for the *Sox1*-*Cre* mouse line. We acknowledge Alex Donovan for technical assistance. We acknowledge the High-Throughput Genomics Group at the Wellcome Trust Centre for Human Genetics (funded by Wellcome Trust grant reference 090532/Z/09/Z) for the generation of the RNA sequencing data and Drs. Brian Nieman and Leigh Spencer Noakes for their MRI sequences. We thank Anthony Graham, Clemens Kiecker and Jeremy Green for critical comments on the manuscript.

---

**Supplementary Fig 1.**
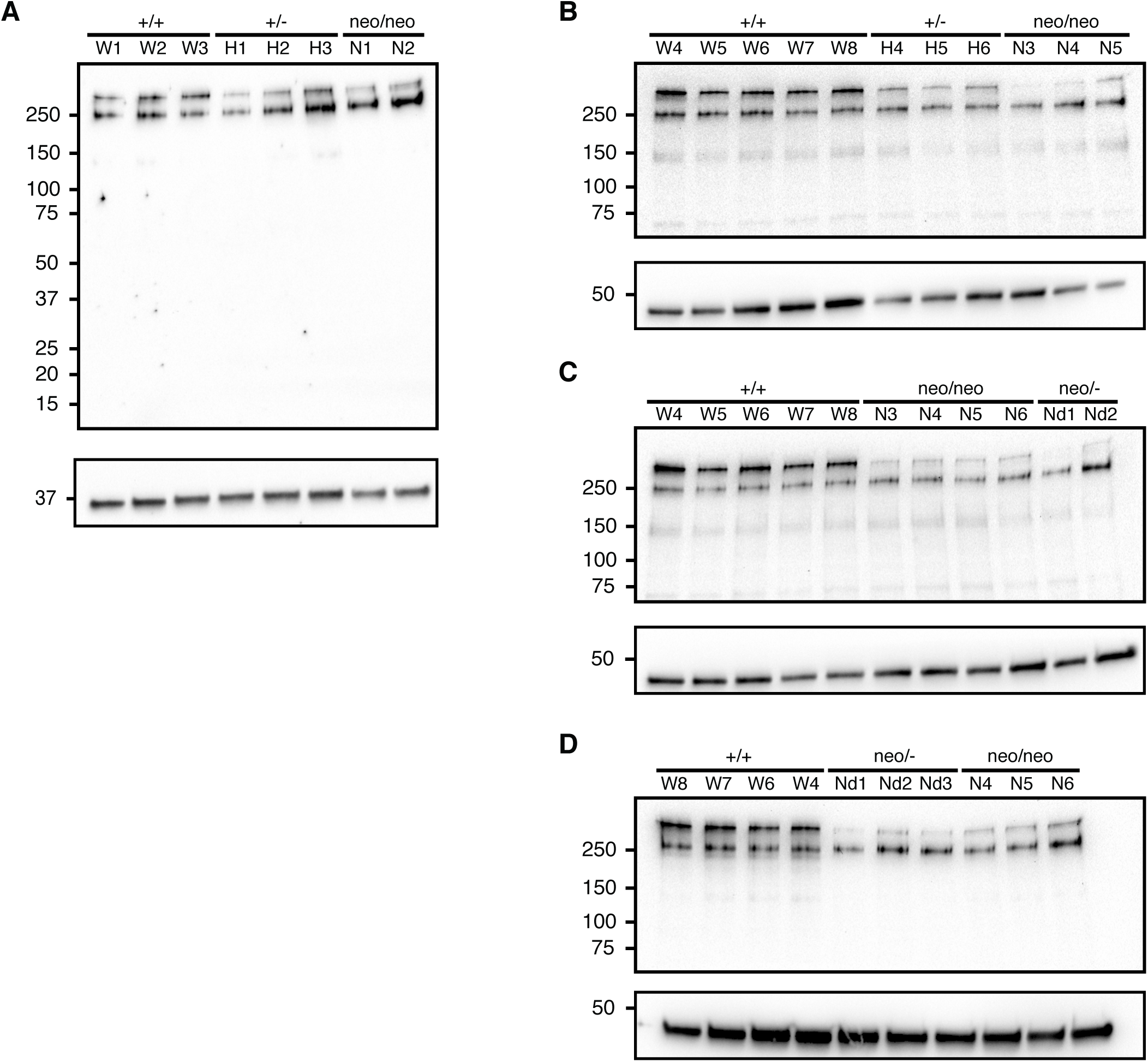
CHD8 expression is reduced in *Chd8*^+/-^, *Chd8^neo/neo^ and Chd8*^*neo/*-^ embryo telencephalic vesicles at E12.5. Whole cell lysate from telencephalic vesicles of indicated genotypes were subjected to western blot analysis using an anti-CHD8 antibody targeting the N-terminal portion of the protein, utilising either GAPDH (A) or β-actin as a loading control (B-D). Labels above individual wells indicate biological replicates used.

**Supplementary Fig 2.**
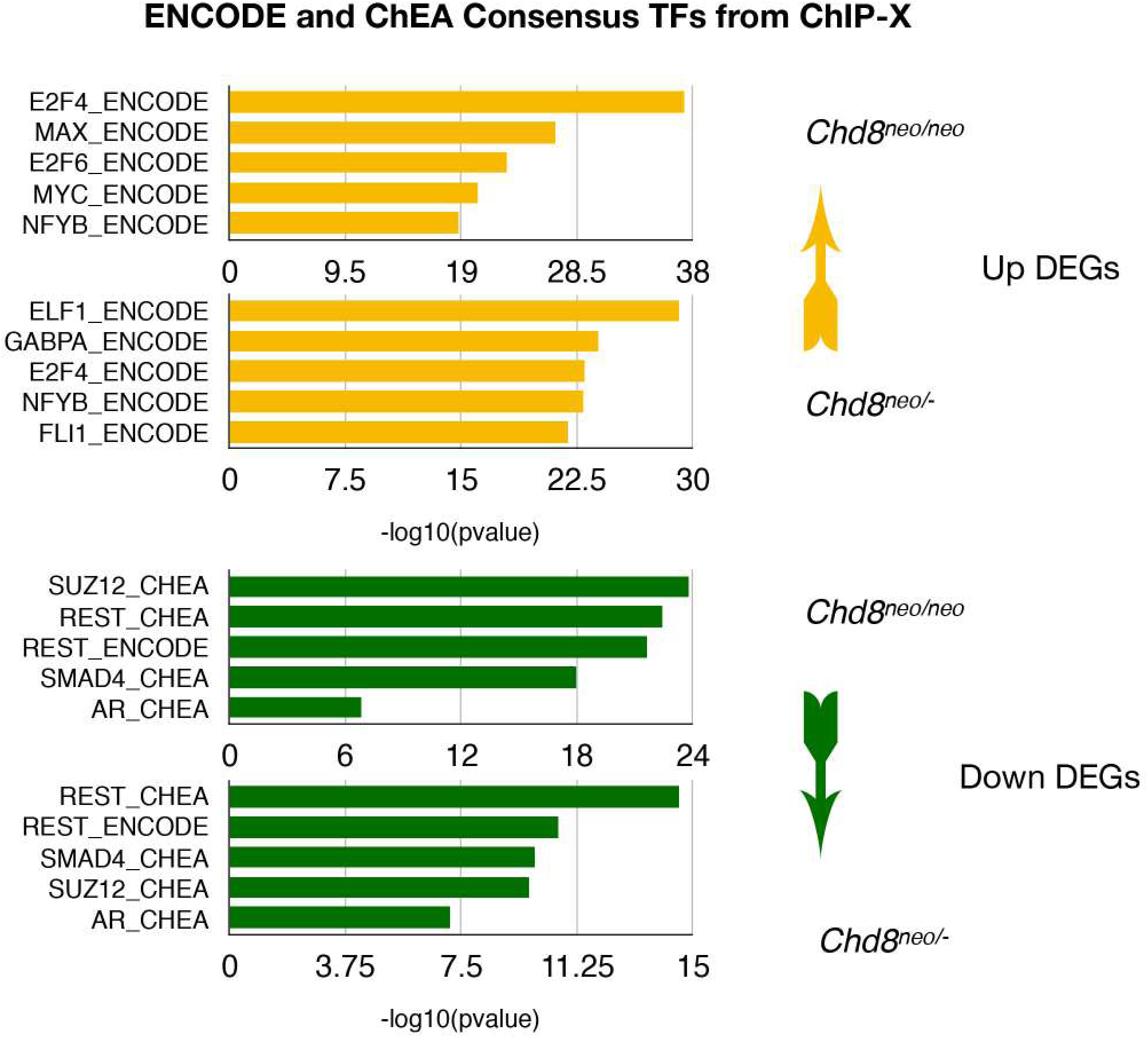
Analysis of transcription factor enrichment using Enrichr. Putative regulatory transcription factors were determined with Enrichr using the “ENCODE and ChEA Consensus TFs from ChIP-X” database with all upregulated DEGs (top yellow panels) and downregulated DEGs (bottom green panels) detected below a 0.05 FDR. The top 5 most significant hits are shown.

**Supplementary Figure 3.**
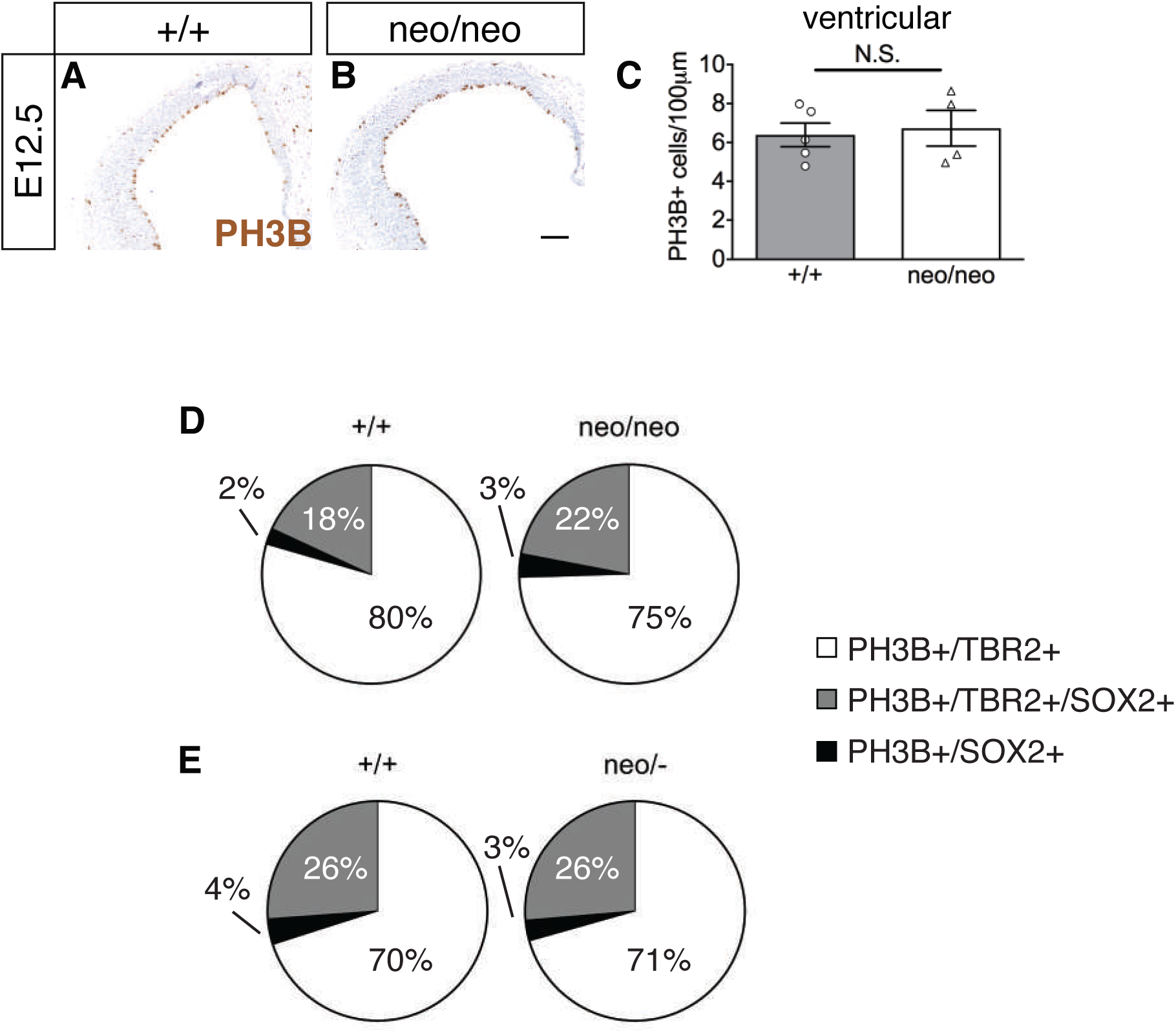
TBR2+ and SOX2+ progenitor proliferation is increased at E14.5 in *Chd8^neo/neo^* embryos. A,B) Immunostaining to detect PH3B+ nuclei (brown) in coronal sections through the telencephalon of E12.5 embryos of indicated genotypes. Scale bar = 100μm. C) Quantification of PH3B+ cells per 100μm of ventricular zone in E12.5 embryos (+/+, n=5; neo/neo, n=4; Mean±SEM, student’s t-test). D,E) Pie charts representing percentages of non-ventricular PH3B+/TBR2+ (white), PH3B+/SOX2+ (grey) and PH3B+/TBR2+/SOX2+ (black) cells out of all non-ventricular PH3B+ cells in E14.5 neo/neo (I) and neo/- (J) embryos compared to their respective wildtype (+/+) littermates.

**Supplementary Figure 4.**
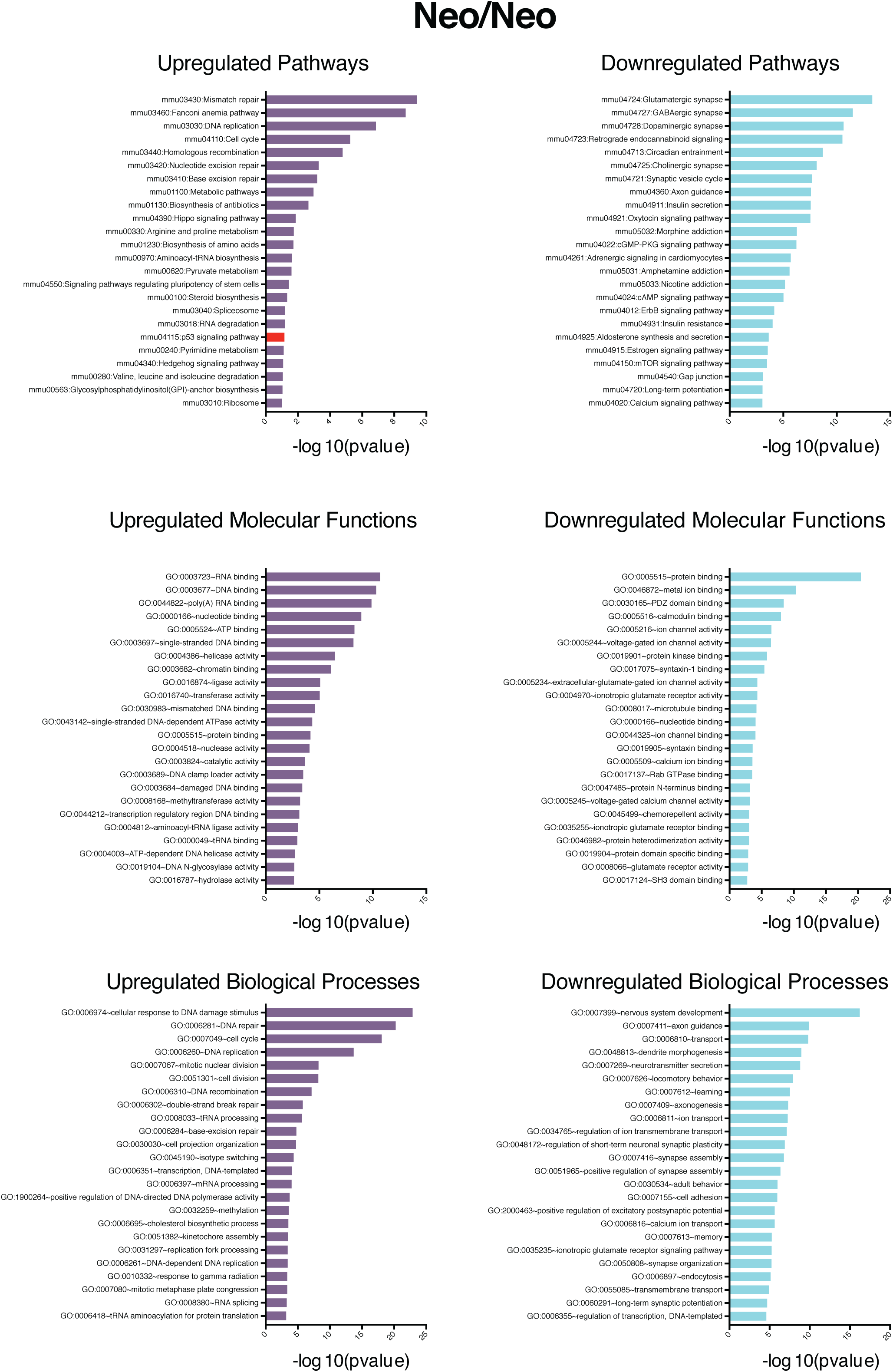
Functional Enrichment Analysis of differentially expressed genes (DEGs) in neo/neo embryos. DEGs (FDR <0.05) were subjected to KEGG pathway enrichment analysis (Top Panels), and screened for Gene Ontology terms in Molecular Function (Middle Panels) and Biological Process (Bottom Panels) categories using the DAVID knowledgebase. In each category, the 25 most significant terms are shown.

**Supplementary Fig 5.**
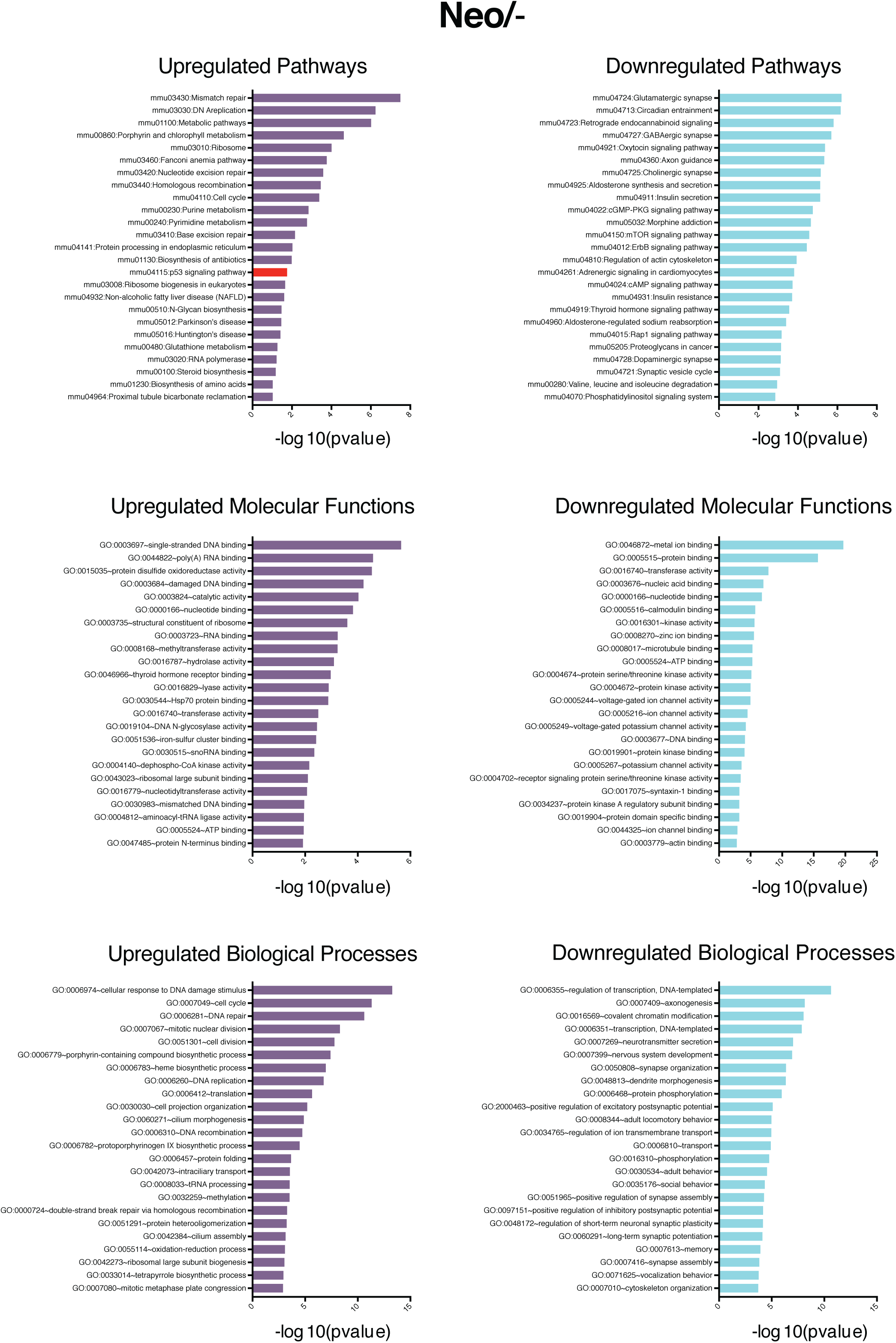
Functional Enrichment Analysis of differentially expressed genes (DEGs) in neo/- embryos. DEGs (FDR <0.05) were subjected to KEGG pathway enrichment analysis (Top Panels), screened for Gene Ontology terms in Molecular Function (Middle Panels) and Biological Process (Bottom Panels) categories using the DAVID knowledgebase. In each category, the 25 most significant terms are shown.

**Supplementary Fig 6.**
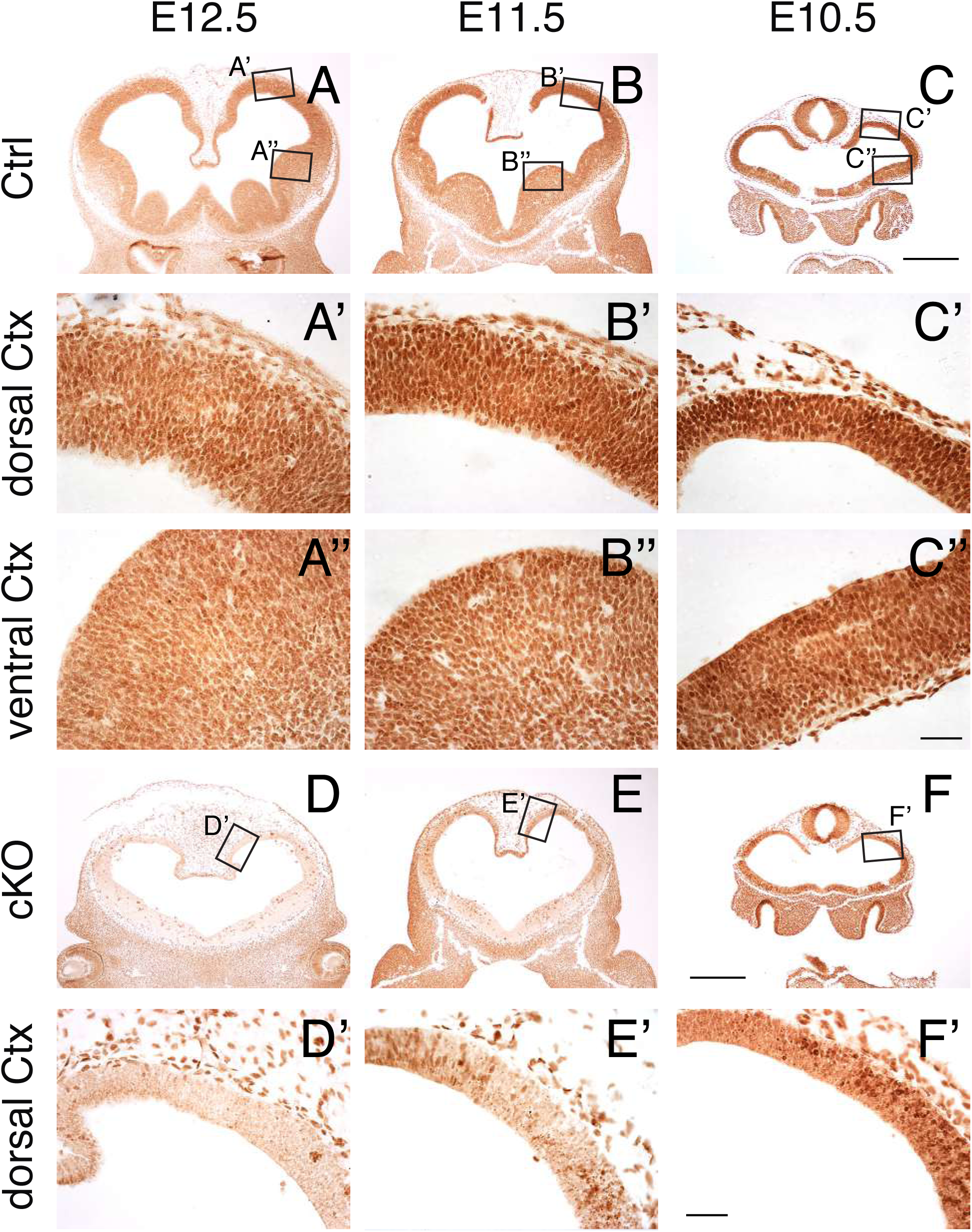
CHD8 is highly expressed during mid-embryonic stages and efficiently removed by recombination with *Sox1*-*Cre*. A-C’’) Immunostaining of E12.5, E11.5 and E10.5 brain sections with an anti-CHD8 antibody. Higher magnification images of dorsal cortex (boxed areas) are shown in A’-C”. Note the presence of nuclear CHD8 protein throughout the pallium and subpallium at E12.5 – E10.5 and higher levels of CHD8 immuno-staining in dorsal (A’-C’) compared to ventral pallium (A’’-C’’). D-F’) Conditional pan-neuronal deletion of *Chd8* results in widespread loss of CHD8 protein in all brain structures. Higher magnification images of dorsal cortex (boxed areas) are shown in D’-F’. Note the loss of CHD8 immunostaining in the neural tube, with unchanged expression in other tissues. Scale bars: A-F: 500μm, A’-F’, A’’-C’’: 50μm

**Supplementary Fig 7.**
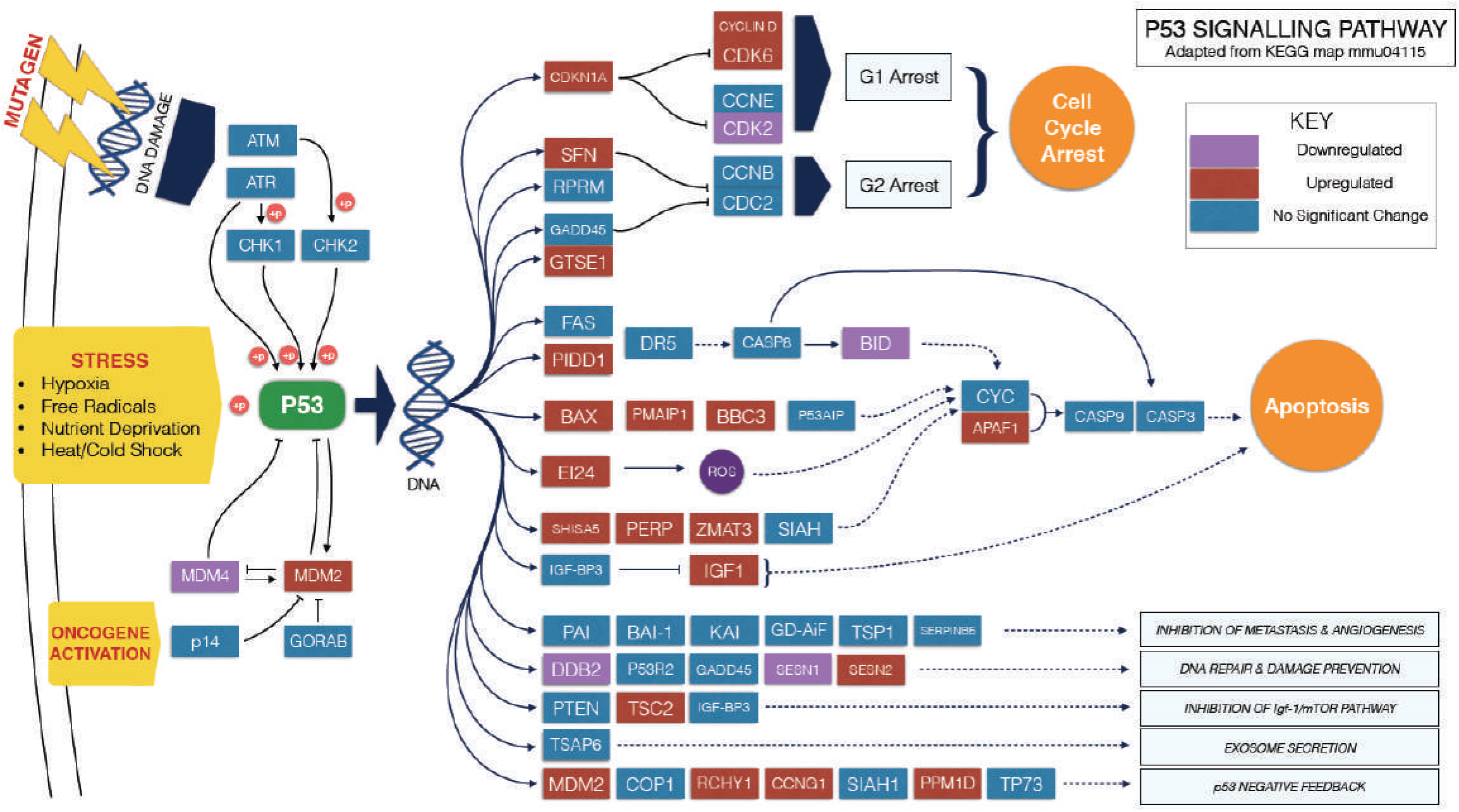
Diagram of the p53 pathway, with differentially expressed genes in the cKO highlighted. KEGG pathway mapping using DAVID of all differentially expressed genes revealed perturbed expression of 27 components of the p53 signalling pathway. Upregulated genes are boxed in red, downregulated genes in purple, and unchanged genes in blue. Solid lines indicate direct interactions whereas broken lines show indirect effects. *Chd8* depletion resulted in more upregulated (22 genes) than downregulated (5 genes) p53 pathway components. The expression of p53 itself (green) was not significantly changed. This figure is adapted from KEGG map mmu04115 (Kanehisa and Goto, 2000).

**Supplementary Table 1**: Raw MRI volumetric data, accompanies Fig. 1D,E.

**Supplementary Table 2**: RNA-seq data from E12.5 neocortices, accompanies Fig. 2.

**Supplementary Table 3**: Gene ontology and pathway analyses of E12.5 RNA-seq data, accompanies Fig. 2I.

**Supplementary Table 4**: RNA-seq and gene ontology analysis of E10.5 cKO telencephalic vesicles, accompanies Fig. 5A,B.

